# Citrus Huanglongbing is an immune-mediated disease that can be treated by mitigating reactive oxygen species triggered cell death of the phloem tissues caused by *Candidatus* Liberibacter asiaticus

**DOI:** 10.1101/2021.08.15.456409

**Authors:** Wenxiu Ma, Zhiqian Pang, Xiaoen Huang, Sheo Shankar Pandey, Jinyun Li, Jin Xu, Diann S. Achor, Fernanda N.C. Vasconcelos, Connor Hendrich, Yixiao Huang, Wenting Wang, Donghwan Lee, Nian Wang

## Abstract

The immune system is critical for keeping animals and plants healthy from pathogens. However, immune-mediated diseases are also common for human. Immune-mediated diseases have not been reported for plants. Here, we present evidence that citrus Huanglongbing (HLB), caused by phloem-colonizing *Candidatus* Liberibacter asiaticus (CLas), is an immune-mediated disease. CLas infection of *Citrus sinensis* stimulated systemic and chronic immune response in the phloem tissues including reactive oxygen species (ROS) production as indicated by H2O2, callose deposition, and induction of immune related genes. Systemic cell death of companion and sieve element cells, but not surrounding parenchyma cells, was observed following ROS production triggered by CLas. ROS production triggered by CLas localized in phloem tissues. The H2O2 concentration in exudates extracted from phloem enriched bark tissue of CLas infected plants reached a threshold of killing citrus protoplast cells, which was suppressed by uric acid (a ROS scavenger) and gibberellin. Foliar spray of HLB positive citrus with antioxidants (uric acid and rutin) and gibberellin significantly reduced both H2O2 concentrations and cell death in phloem tissues induced by CLas and reduced HLB symptoms. RNA-seq analyses of CLas infected and health *C. sinensis* support that CLas causes oxidative stress. In sum, HLB is an immune-mediated disease and both mitigating ROS via antioxidants and promoting plant growth can reduce cell death of the phloem tissues caused by CLas, thus controlling HLB.

---

Both plants and animals utilize the immune system to fight off pathogens. However, some human diseases are mediated by the immune response. Immune-mediated diseases include diseases due to inflammation stemming from the immune response to certain microbes and environmental antigens ^1^. For example, inflammatory bowel disease is disorders that involve chronic inflammation of the digestive tract. Bacterial pathogens can instigate chronic inflammation that leads to diseases beyond the damaging effect of pathogenicity factors ^2^. In addition, autoimmune diseases, such as allergic diseases, and allergic asthma are also immune mediated diseases.

The damaging effect of plant diseases has been assumed to directly result from the impact of pathogenicity factors of corresponding pathogens ^3^. Common pathogenicity factors include effectors, toxins, cell wall degrading enzymes, and biofilm that are directly responsible for causing disease symptoms. For instance, the transcriptional activator-like effector PthA4 is responsible for the hypertrophy and hyperplasia symptoms of citrus canker caused by *Xanthomonas citri* subsp. citri ^4^. Xylem blockage caused by biofilm of *Xylella fastidiosa* is known to lead to the wilting of grapevine plants with Pierce’s disease ^5^. Citrus Huanglongbing (HLB, also known as citrus greening) is currently the most devastating citrus disease and causes billions of dollars economic losses worldwide annually. HLB is caused by the phloem-colonizing *Candidatus* Liberibacter asiaticus (CLas), *Ca*. L. americanus and *Ca*. L. africanus that are vectored by either Asian citrus psyllid (*Diaphorina citri*) or African citrus psyllid (*Trioza erytreae*)^6^. Among them, CLas is the most prevalent worldwide.

Despite its economic importance, how *Ca*. Liberibacter causes damages to the infected citrus plants remains poorly understood. One reason for such a delay is that HLB pathogens have not been cultured in artificial media. No pathogenicity factors have been confirmed to be responsible for the HLB symptoms including the characteristic blotchy mottle on leaves, hardened and upright small leaves, stunt growth, and root decay ^6^. Here we present evidence that citrus HLB is an immune-mediated disease. This hypothesis explains most phenomena observed for HLB, fits the genetic information of *Ca*. Liberibacter spp., and also provides guidance regarding HLB management.

## CLas does not contain pathogenicity factors that directly cause HLB symptoms

We conducted a comprehensive analysis of CLas proteins and did not identify any homologs with known pathogenicity factors that are directly responsible for causing disease symptoms. *Ca*. Liberibacter spp. do not contain the type II, III, and IV secretion systems that secrete such pathogenicity factors. To test whether CLas contains pathogenicity factors responsible for causing HLB symptoms, 47 predicated virulence factors including serralysin and hemolysin (substrates of type I secretion system) and proteins containing Sec secretion signals (Table S1) ^7^ were overexpressed in *Arabidopsis thaliana*, *Nicotiana tabacum* or *Citrus paradisi*. None of the overexpressed CLas proteins caused HLB-like symptoms, consistent with the bioinformatic analyses that CLas does not contain pathogenicity factors that directly cause HLB symptoms. Intriguingly, multiple characterized proteins of CLas, such as SDE1, SDE15, and SahA, suppress plant immune response ^8–10^, suggesting that CLas triggers immune response, which, we hypothesize, is responsible for causing the devastating damages of the HLB disease, mimicking the immune-mediated diseases of human.

## CLas infection triggers immune response and cell death in the phloem tissues

Next, we tested whether and how CLas triggers immune response and cell death. Newly emerged citrus flush from HLB positive citrus trees is free of CLas for a short period of time. We trimmed HLB positive and healthy two-year-old *C. sinensis* ‘Valencia’ trees in a greenhouse to trigger young flush and conducted dynamic analyses of the relationship between CLas infection, immune response, cell death, and HLB symptom development. CLas was detected in young leaves of HLB positive trees at approximately 15 days post bud initiation based on quantitative PCR (qPCR) (Fig. 1). The H2O2 (an indicator of reactive oxygen species (ROS)) content in CLas positive flush was significantly higher than that of healthy plants at 15- and 21-day post bud initiation (Fig. 1A). Significantly more callose deposition, an indicator of immune response ^11^, was observed in CLas positive flush than that of the healthy plants starting at 18 days post bud initiation and thereafter (Fig. 1B). On the other hand, significantly more starch accumulation was observed in CLas positive samples than in healthy samples starting at 18 days post bud initiation and afterwards (Fig. 1C). Symptoms began to appear at approximately 40 days post bud initiation. It appears that CLas infection triggered plant immune response, such as ROS (e.g., H2O2) production and callose deposition, followed by symptom development.

**Fig. 1.**
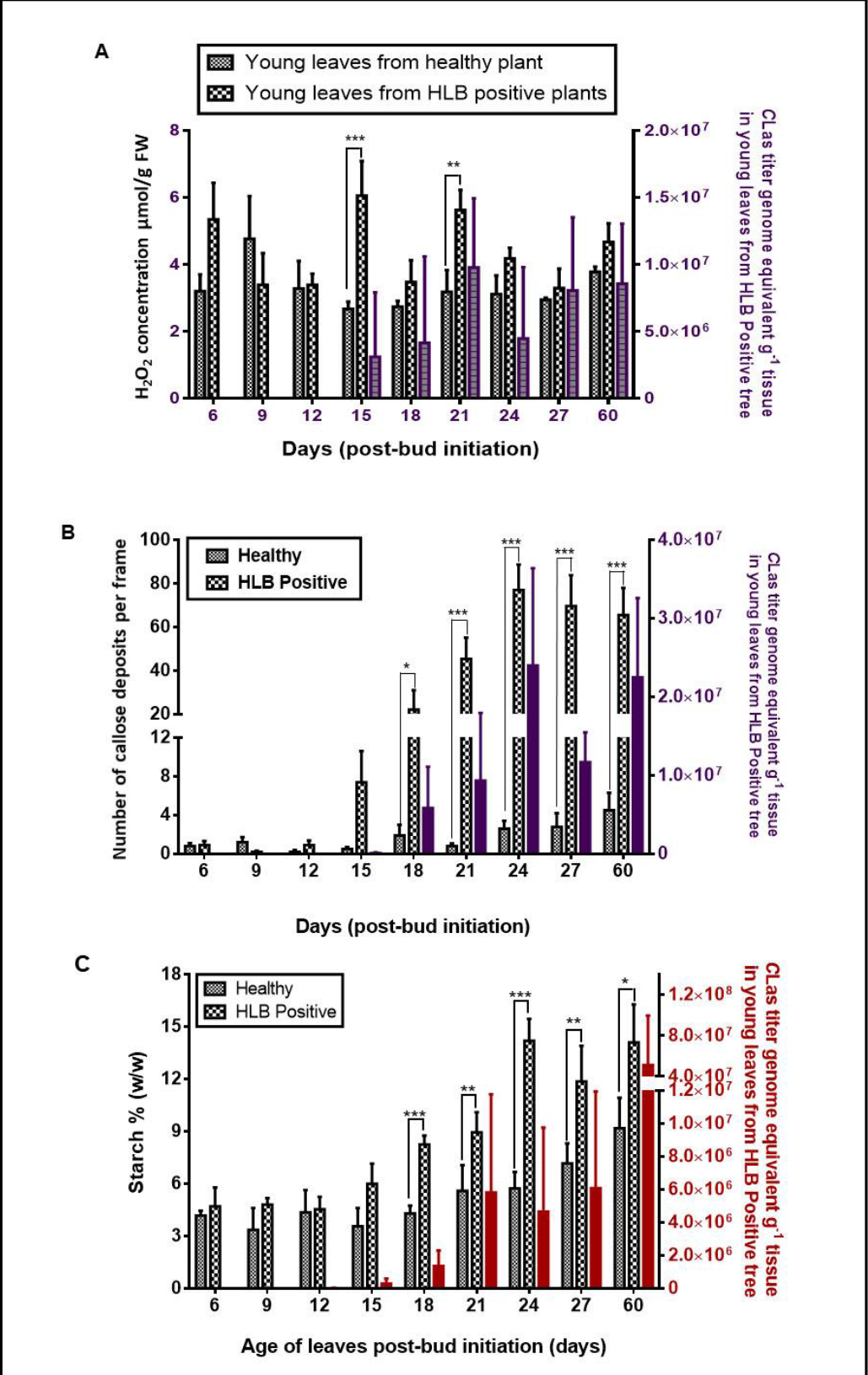
CLas infection causes ROS production, callose deposition, and starch accumulation in young citrus leaves. (A) Quantification of H2O2 in young leaves of two-year-old HLB positive and healthy *C. sinensis* ‘Valencia’ trees. n=4. (B) Quantification of callose depositions of young leaves by staining with aniline blue and observed under an epifluorescence microscope. Data shown are mean ± SD (standard deviation), n=10. Callose depositions were counted from ten leaves from five trees. (C) Starch content in young flush. Data shown are mean ± SD; n=4. HLB positive and healthy two-year-old *C. sinensis* ‘Valencia’ trees were trimmed to trigger young flush in the greenhouse. qPCR was conducted to quantify CLas titers in newly emerged leaves at different time points after bud initiation. Data shown are mean ± SD; n=4. Experiment was repeated twice with similar results. *, **, and *** indicate *P* value < 0.05, 0.01, and 0.001, respectively, compared with healthy young leaves and calculated using Student’s t-test.

To have a better understanding of the nature of the immune response induced by CLas, we conducted temporal expression analyses of immune marker genes (PR1, PR2, PR3, and PR5) in young leaves at 15-, 18-, 21-, 24-, 27-, and 60-days post-bud initiation for the CLas infected and healthy *C. sinensis* plants. PR genes were consistently induced by CLas despite some fluctuations between 15- and 60-day-post-bud initiation (fig. S1).

We observed cell death of sieve element and companion cells via transmission electron microscopy (TEM) analysis of asymptomatic young leaves of HLB positive *C. sinensis* ‘Valencia’ trees (Fig. 2A-D), indicating cell death of phloem tissues occurs prior to the appearance of HLB symptoms. CLas was observed in phloem tissues with intact sieve element and companion cells, but not in sieve element cells undergoing the cell death process (Fig. 2C-E). Particularly, we observed some sieve element and companion cells undergoing cell death while others remained intact in the same field (Fig.2D), explaining the reduced function of the phloem tissues, rather than a complete loss of function. This accords with the observation that CLas moves primarily vertically, but not laterally during infection ^12^. TEM observation of CLas infected leaves confirmed that cell death was limited to sieve element and companion cells, but not occurring in surrounding parenchyma cells (Fig. 2C).

**Fig. 2.**
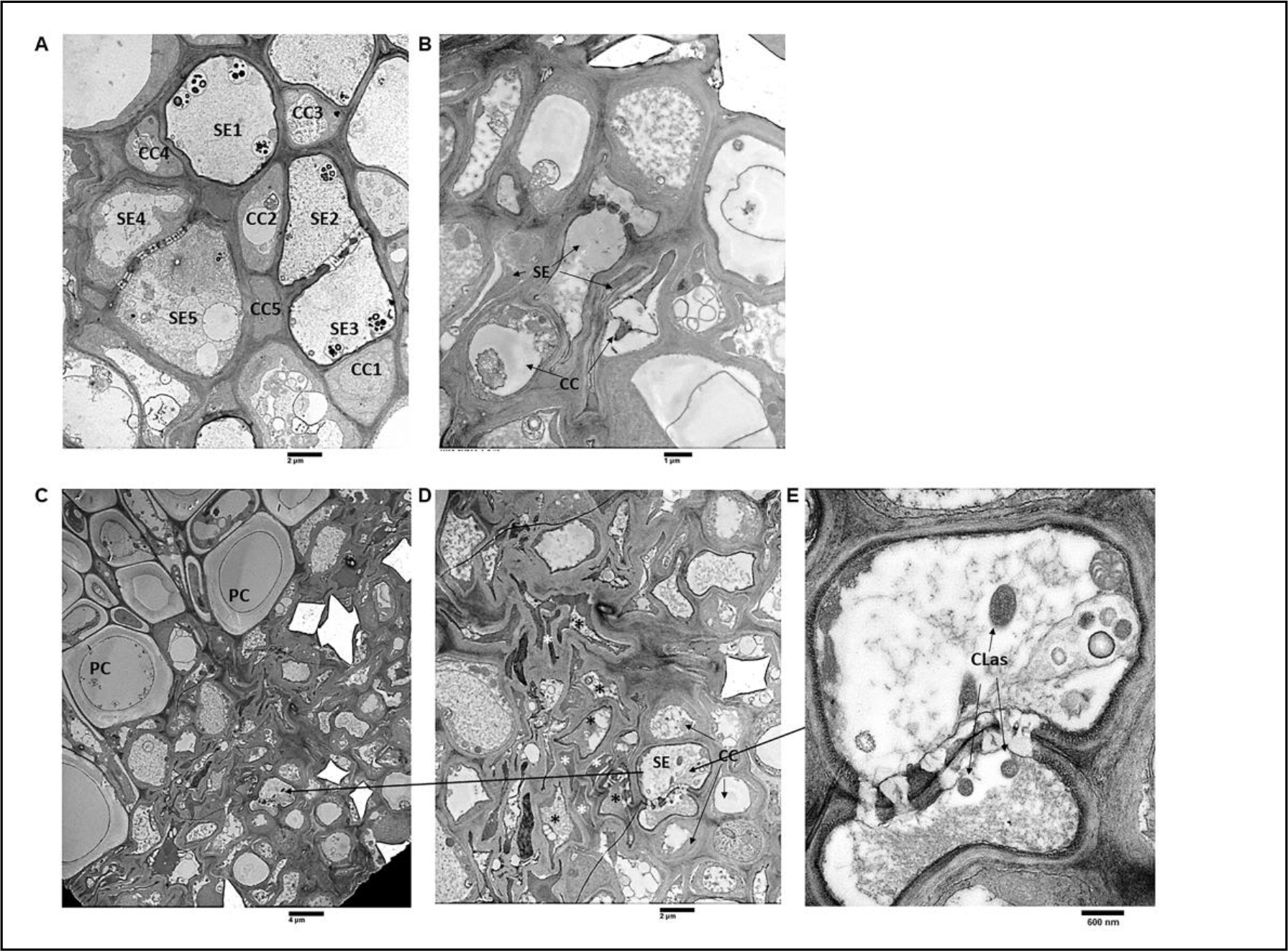
Transmission Electron Microscopy (TEM) analysis of asymptomatic young leaves of CLas infected *Citrus sinensis*. (A) Heathy leaf of *C. sinensis* ‘Valencia’. CC (companion cells): typical healthy companion cells with large central nucleus, dense cytoplasm, numerous mitochondria. SE (sieve element 1-3): typical sieve elements showing parietal “s” plastids, centrally distributed p protein, and lateral sieve plate showing minimum callose. SE (4-5): developing sieve elements showing plastids with no starch, intact central vacuole and non-dispersed phloem protein. (B) Asymptomatic young leaf of CLas infected *C. sinensis*. CC undergoing cell death. (C, D, and E) Asymptomatic young leaves showing cell death of sieve element cells and companion cells, but not parenchyma cells (PC). Black * indicates cell death of sieve element cells. White * indicates cell death of companion cells. (D) Enlarged section of C showing both intact sieve element and companion cells as well as sieve element and companion cells undergoing cell death. (E) Enlarged section of D showing intact sieve element cells containing CLas.

We confirmed the cell death in *C. sinensis* mature leaves showing different symptoms based on trypan blue staining. No cell death was observed in healthy leaves collected from CLas-free plants. Cell death was observed in CLas positive asymptomatic leaves, and leaves with mild or severe HLB symptoms and correlated positively with symptom development (Fig. 3A). In addition, cell death was observed along the vascular tissues, congruent with the TEM observation of the death of sieve element and companion cells in the midribs of CLas infected mature leaves and CLas positive stem tissues (figs. S2 and S3). More cell death was observed with increasing CLas titers, suggesting CLas infection is responsible for the cell death of the phloem tissues (figs. 2 and 3). TEM observation demonstrated that cell death of companion and sieve element cells occurred concurrently without obvious difference in timing, in agreement with their function together as a unit ^13^. We observed significantly higher H2O2 concentrations in CLas infected mature leaves than CLas free leaves (Fig. 3B). In addition, significantly higher H2O2 concentrations were detected in the exudates of phloem enriched tissues of CLas positive stems than that of stems of CLas-free trees (Fig. 3C).

**Fig. 3.**
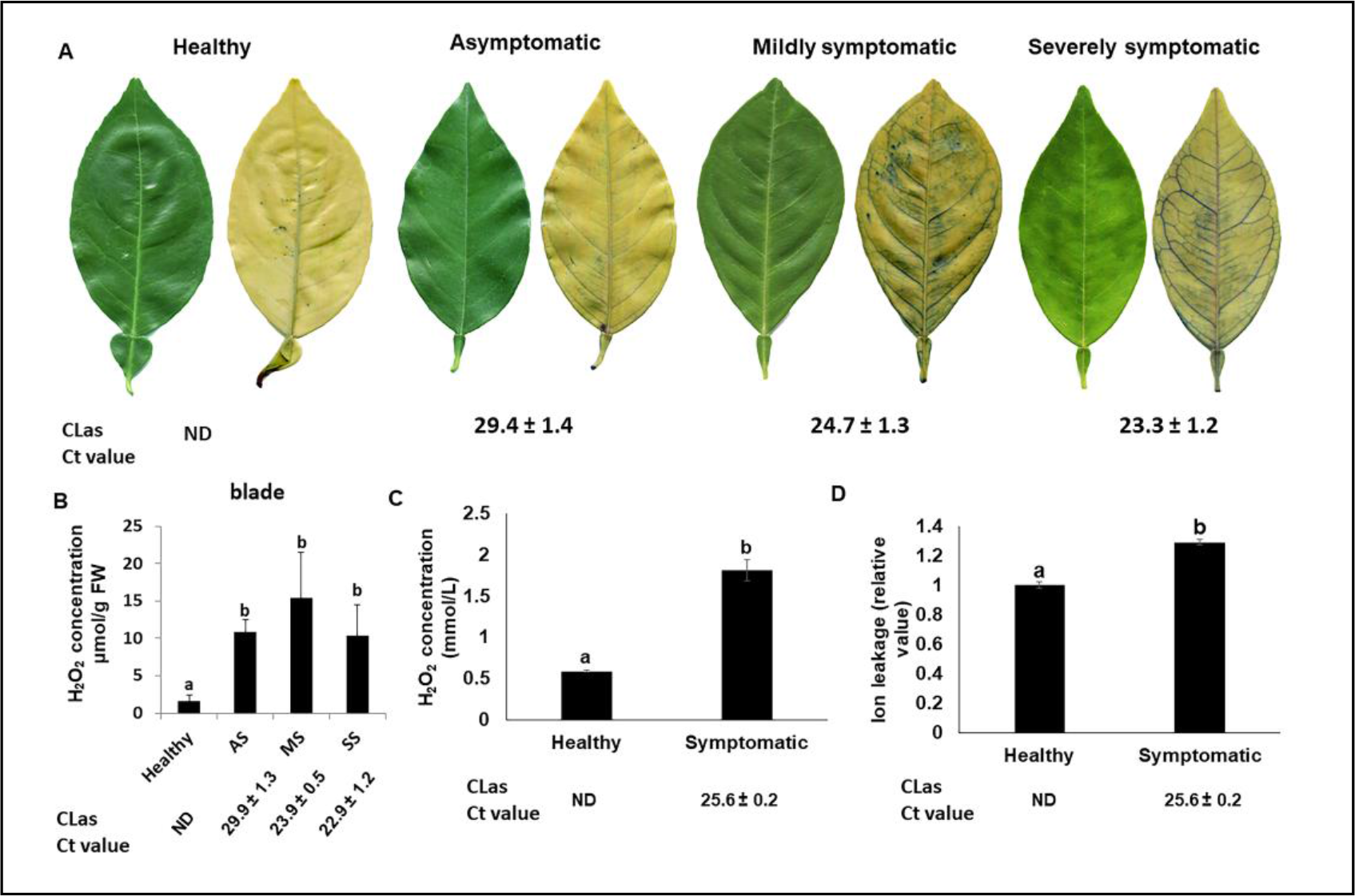
C*a*ndidatus Liberibacter asiaticus (CLas) induces ROS production and cell death in phloem tissues of *C. sinensis*. (A) Trypan blue staining assay to detect cell death in leaves of CLas infected *C. sinensis* ‘Valencia’ trees. (B) Determination of H2O2 concentration in CLas negative or positive leaves showing different symptoms. Mean and SD are shown. (C and D) H2O2 (C) and ion leakage (D) assays of exudates extracted from phloem enriched bark tissues. Statistical differences were analyzed using one way ANOVA with Bonferroni Correction (*P*<0.05) for B and using Student’s t-test for C and D. Different letters above the columns indicate statistical differences (*P*<0.05). ND: non-detected. For A and B: AS: asymptomatic. MS: Mildly symptomatic. SS: Severely symptomatic. AS, MS, and SS were collected from HLB-positive trees. For C and D, H: stems of spring sprouts were collected from healthy branches of CLas-negative plants. S: stems of spring sprouts were collected from branches with HLB symptomatic leaves of CLas-positive plants. Each experiment contains four biological replicates. The experiments were repeated at least twice with similar results. Ct values of CLas of the tested samples were indicated.

Cell death is usually accompanied by ion leakage. Surprisingly, no difference was observed in ion leakage between leaf blades or midribs of healthy, asymptomatic, mildly symptomatic, and severely symptomatic leaves (fig. S4), contrary to the TEM and trypan blue staining data (Figs. 2 and 3A, and fig. S2). However, the ion leakage values of the exudates extracted from phloem enriched bark tissues of CLas infected samples were significantly higher than that of healthy samples (Fig. 3D), consistent with that cell death happens in the phloem tissues, but not in surrounding parenchyma cells (Fig. 2C). The negative ion leakage data related to leaf blades and midribs of CLas infected samples (fig. S4) probably result from the mask effect of parenchyma cells because companion and sieve element cells make up only approximately 1% of the total cell population in plants ^14^.

Next, we used callose deposition as an indicator to investigate the localization of the immune response in citrus leaves in response to CLas infection. For this test, we investigated callose deposition in different sections of asymptomatic and symptomatic leaves of HLB-positive *C. sinensis* trees (Fig. 4A-G). The callose deposition in the petiole, midrib, and lamina of asymptomatic leaves was significantly lower than that in their counterparts of symptomatic leaves (Fig. 4, A-G). No callose deposition was observed in the CLas-free lamina of asymptomatic leaves (Fig. 4C). Collectedly, the correlation between callose deposition and CLas titers suggests that CLas is responsible for inducing callose deposition (Fig. 4A-F) as observed in the past ^15^. Callose deposition was observed only in the phloem tissues as observed previously ^16^, but not in mesophyll cells. In contrast, for pathogens infecting the apoplast, such as *Xanthomonas*, callose is deposited between the plasma membrane and the cell wall at the site of pathogen attack ^17, 18^. Hence, CLas induces systemic immune response in the phloem tissues following systemic CLas infection.

**Fig. 4.**
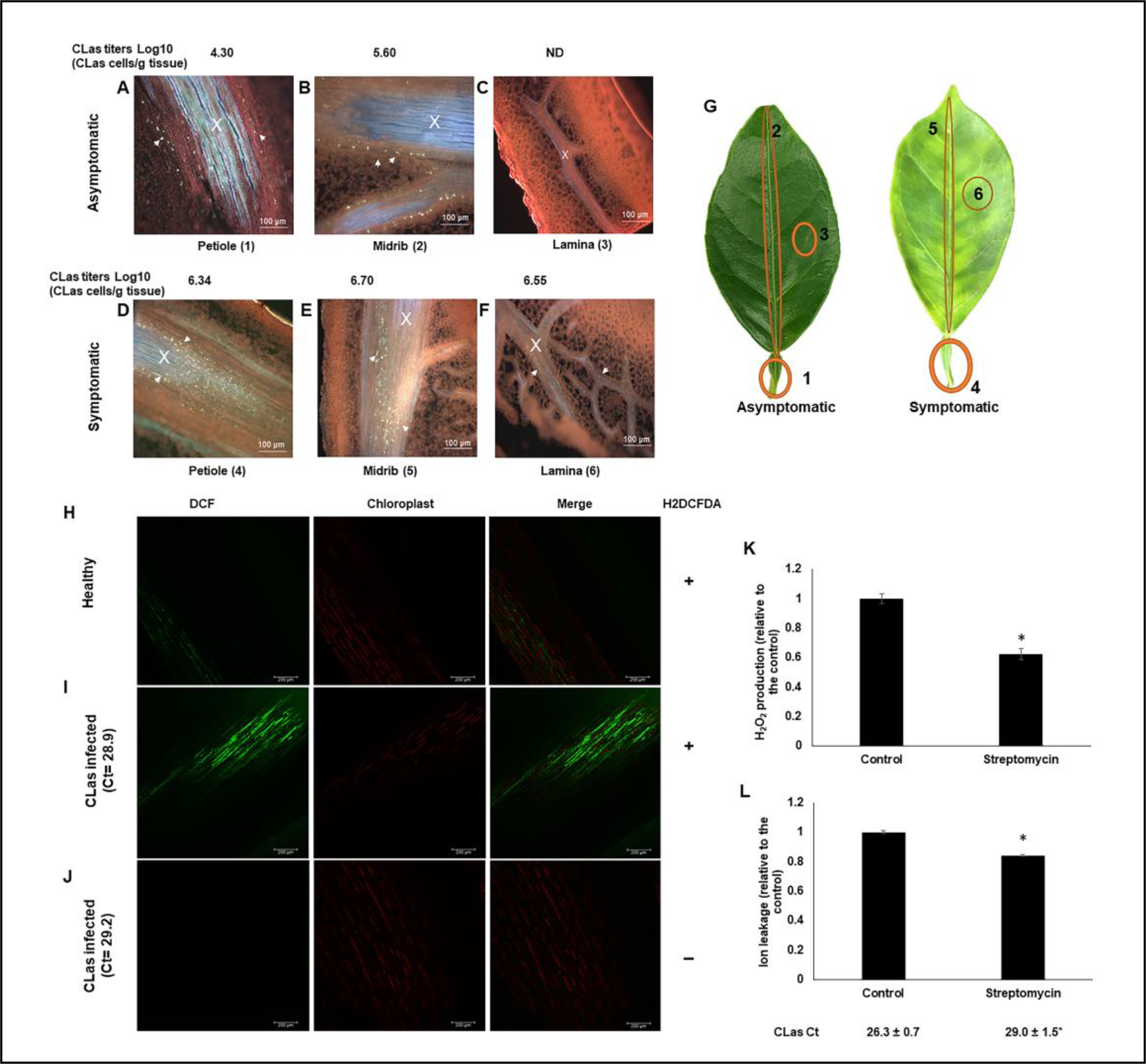
CLas infection induces callose deposition, H2O2 production and cell death in the phloem tissues of *C. sinensis*. (A-F) *C. sinensis* ‘Hamlin’ leaf samples were fixed with FAA solution overnight, sectioned and stained with 0.005% aniline blue solution prior to analysis. Xylem is marked with an X, and callose deposition is indicated with arrowheads. Asymptomatic samples: (A) Petiole, (B) Midrib, (C) Lamina; Symptomatic samples: (D) Petiole, (E) Midrib, and (F) Lamina. Pictures are representatives of 12 replicates. ND: Non-detected. (G) schematic representation of samples used for callose deposition assays (A-F). CLas titers for each section were quantified by qPCR. (H-J) H2O2 production in the phloem tissues. Healthy (H) and CLas infected (I) *C. sinensis* ‘Valencia’ bark tissues visualized with 2’,7’- dichlorodihydrofluorescein diacetate (H2DCFDA) under a confocal laser microscope. CLas infected (J) *C. sinensis* ‘Valencia’ bark without H2DCFDA was used as a control. “+” indicates with H2DCFDA treatment. “-” indicates without H2DCFDA treatment. (K-L) Effect of killing CLas with streptomycin on H2O2 concentration (K) and cell death (L) in the phloem tissues. CLas positive five-year-old sweet orange trees were trunk-injected with streptomycin. Non-treatment was used as the negative control. The tests were conducted 7 days after trunk injection of streptomycin. Four biological replicates were used. * indicates significant statistical difference (*P*<0.05) based on Student’s t-test.

To further verify that CLas induces immune response in the phloem tissues, we monitored ROS formation and localization in phloem tissues using the fluorescent probe 2’,7’- dichlorodihydrofluorescein diacetate (H2DCFDA) and confocal laser microscopy. H2DCFDA is a commonly used cell-permeable probe for measuring cellular H2O2 ^19^. Significantly higher H2O2 was detected in the phloem tissues of CLas infected citrus plants than that of CLas-free plants (Fig. 4H-J).

Next, we aimed to further establish the causal relationship between CLas infection and ROS induction and cell death in phloem tissues. For this assay, CLas-positive 5-year-old *C. sinensis* trees were treated with streptomycin to kill CLas via trunk injection ^20^. At 7 days post treatment, streptomycin significantly reduced CLas titers, H2O2 content and ion leakage in the phloem tissues (Fig. 4K-L). Collectively, we have established the causative relationship between CLas infection and ROS induction and cell death in the phloem tissues.

## HLB caused cell death is instigated by ROS

It is known that cell death can be initiated by ROS ^21^. At high concentrations, ROS triggers necrotic cell death, but induces programmed cell death below the ROS threshold ^22^. Next, we analyzed H2O2 contents in *C. sinensis* leaves triggered by CLas infection. The H2O2 concentration induced by CLas infection in young leaves was approximately 6 µmol g^−1^ FW, but reached 10-15 µmol g^−1^ FW in mature leaves (Fig. 3B). Moreover, this method probably underestimated the H2O2 concentration in the phloem tissues because it could not differentiate phloem cells, where H2O2 concentrates (Fig. 4H-J), from parenchyma cells. Intriguingly, *Xanthomonas citri* subsp. citri, another bacterial pathogen of citrus, infection of kumquat (*Citrus japonica*, syn: *Fortunella crassifolia*) triggers H2O2 production at 2 days after inoculation which peaks (9.86 µmol g^−1^ FW) at 8 days after inoculation, eventually leading to cell death ^23, 24^. *X. citri* subsp. citri induced cell death is a slow process and happens at approximately 6-8 days after inoculation ^24^. Hence, we conjectured that ROS induced by CLas reaches the threshold necessary to trigger the death of companion and sieve element cells in mature leaves, but possibly not in the early stage of infection of young leaves before CLas titers reach a certain threshold. The ROS production triggered by CLas is distinct from that triggered by incompatible pathogens, which is typified by a biphasic oxidative burst ^25^. Instead, the ROS production triggered by CLas is chronic, and was observed in young leaves during early infection stages as well as in CLas infected mature leaves consistently, probably triggered by CLas colonizing and multiplying in the previously unoccupied phloem tissues.

Furthermore, the H2O2 concentrations in the exudates extracted from the phloem enriched bark tissues from symptomatic (1.80 ± 0.13 mmol/L) CLas positive branches were significantly higher than that (0.59 ± 0.01 mmol/L) of healthy trees (Fig. 3C). H2O2 induces necrosis of immortalized rat embryo fibroblasts at a concentration of 700 μmol/L ^26^. H2O2 at concentrations of 1.8 mmol/L but not 0. 6 mmol/L or lower induced cell death of *C. sinensis* protoplast cells (Fig. 5A and B; fig. S5). Addition of uric acid (0.2 mM), a ROS scavenger, reduced both H2O2 concentration (Fig. 5C) and cell death (Figs. 5, A and B), indicating H2O2 induced by CLas alone can cause cell death of phloem tissues. It is important to note that growth of plants (e.g., *Arabidopsis*) is inhibited by 1 mM H2O2 ^27^, partly explaining the growth stunting phenotype of CLas infected young trees.

**Fig. 5.**
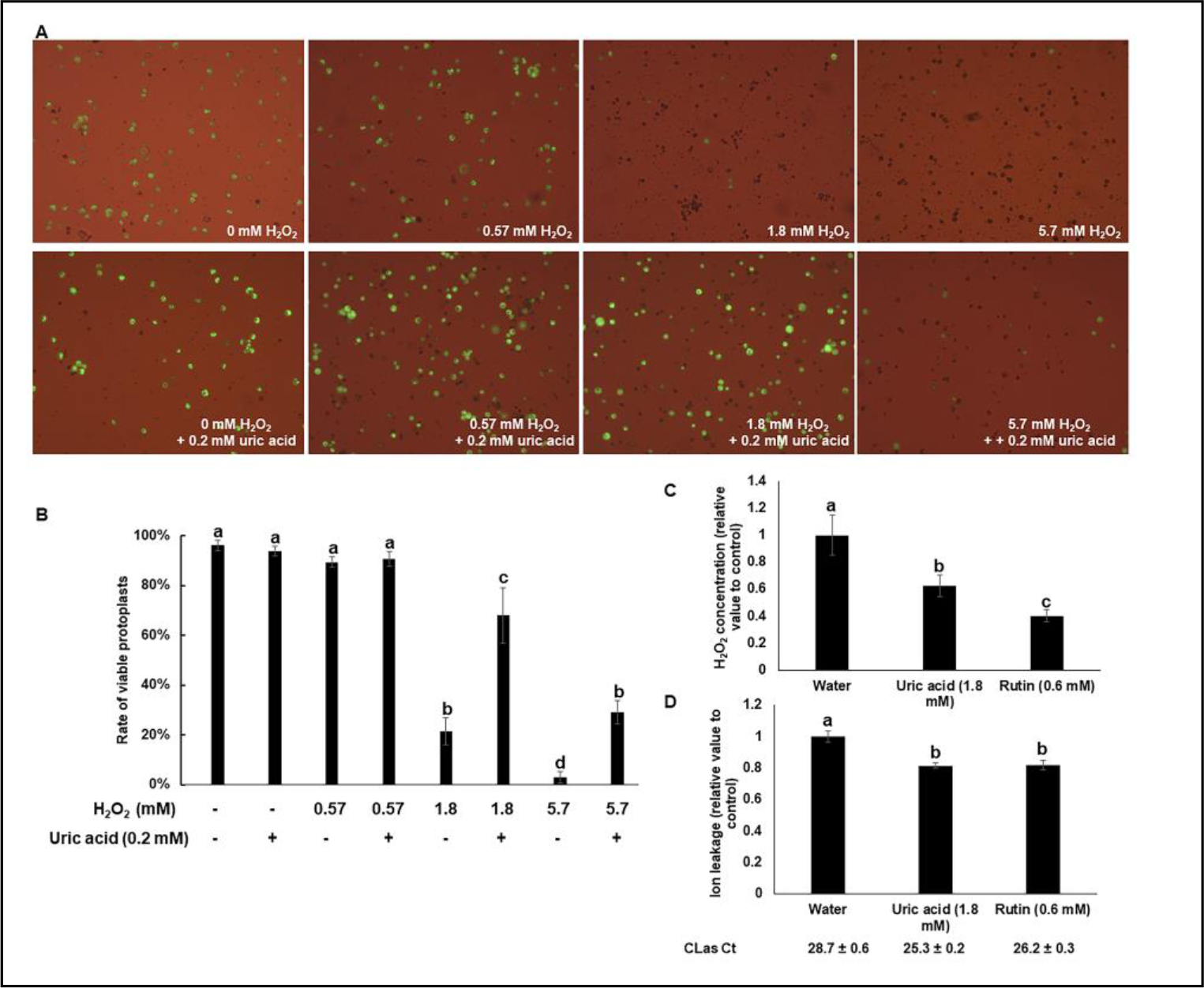
ROS is responsible for cell death of the phloem tissues of CLas infected citrus. (A and B) H2O2 kills protoplast cells of *C. sinensis*. (A) Freshly prepared protoplast cells of *C. sinensis* were treated with different concentrations of H2O2 with or without the antioxidant uric acid (0.2 mM) for 24 h and tested for viability via fluorescein diacetate (FDA) staining. (B) Quantification of viable protoplast cells in different treatments of A. (C and D) HLB-positive plants were treated with antioxidants via foliar spay weekly for six weeks. The exudates extracted from phloem enriched bark tissues were used for detection of H2O2 and ion leakage. (C) Uric acid (1.8 mM) and rutin (0.6 mM) reduce H2O2 concentration triggered by CLas in the phloem tissue. (D) Uric acid (1.8 mM) and rutin (0.6 mM) reduce ion leakages in the phloem tissues infected by CLas. Statistical differences were analyzed using one way ANOVA with Bonferroni Correction (*P*<0.05). Each experiment contains four biological replicates. Different letters above the columns indicate statistical differences (*P*<0.05). Ct values of CLas of the tested samples were indicated.

In addition to H2O2, ROS induced by pathogens include hydroxyl radicals, superoxide anions, and singlet oxygen. To further corroborate that HLB caused cell death is instigated by ROS, we conducted weekly foliar spray of HLB positive *C. sinensis* ‘Valencia’ trees with antioxidants uric acid (1.8 mM) and rutin (0.6 mM). Six weeks later, analysis of the exudates extracted from the phloem-enriched bark tissues demonstrated that both uric acid and rutin treatments reduced both ROS production, as indicated by H2O2 concentration (Fig. 5C), and cell death (Fig. 5D). Taken together, CLas infection of citrus phloem tissues induces ROS production, which subsequently causes cell death of phloem tissues.

## CLas infection significantly affects pathways related to oxidative stress and immune responses

Next, we investigated the gene expression profiles of *C. sinensis* in response to CLas infection that were conducted previously, comprising 9 studies including different tissues (leaf, stem, and fruit), different environments (greenhouse or groves), and different infection stages (Table S2). Enrichment analyses of differentially expressed genes (DEGs) clearly demonstrated that the expression of genes related to ROS and immune response are significantly affected by CLas infection (Table S3). The combined analysis showed an overall downregulation of antioxidant enzymes and upregulation of transmembrane localized NADPH oxidases, known as RBOHs, explaining the oxidative stress response in response to CLas infection (fig. S6). Critically, expression of respiratory burst oxidative homolog D (RbohD) genes, which encode an enzyme implicated in the generation of ROS during the defense response, was induced by CLas infection in most occasions. RBOHD is a main producer of ROS upon PAMP recognition and required for cell death initiated after pathogen detection ^28^. The combined analysis implies that CLas infection causes oxidative stress to citrus in most conditions, consistent with H2O2 production triggered by CLas (Figs. 1A, 3B-C, 4H-K, and 5C). The combined analysis also revealed complex expression changes associated with the immune response pathways in response to CLas infection (fig. S7). Intriguingly, approximately 66 NLR genes showed overall induction by CLas in multiple studies (fig. S7B). In sum, transcriptome analyses of sweet orange in response to CLas infection support our hypothesis that HLB is an immune-mediated disease that results from ROS induced cell death of phloem tissues triggered by CLas.

## Suppressing ROS mediated cell death mitigates HLB symptoms

Antioxidants, and immunoregulators are commonly used to treat human immune-mediated diseases by halting or reducing ROS mediated cell death ^29–31^. Correspondingly, we tested whether growth hormones gibberellin (GA), and antioxidants (uric acid and rutin) mitigate ROS mediated cell death triggered by CLas infection, thus blocking or reducing HLB symptoms. GA is selected because it is a known plant growth hormone and modulates PAMP-triggered immunity and PAMP-induced plant growth inhibition ^32^. Both uric acid and rutin are well-known ROS scavengers ^33, 34^.

Foliar spray of HLB positive *C. sinensis* trees with GA at both 5 mg/L and 25 mg/L reduced H2O2 and ion leakage caused by CLas (Fig. 6, A and B). Consistent with the foliar spray results, GA (5 mg/L) also suppressed death of *C. sinensis* protoplast cells caused by 1.8 mM H2O2 (Fig. 6, C and D). Six weeks after foliar spray, the treated plants showed reduced HLB symptoms compared with that before treatment, whereas plants with water treatment developed more severe HLB symptoms in the same period (fig. S8). In addition, it was obvious that GA promoted growth as indicated by new flushes, which were not observed for the water control in the same duration (fig. S8). GA is registered to be used on citrus in the US, which enables large scale field trials to test its effect against HLB. Foliar spray of *C. sinensis* with GA significantly reduced HLB disease symptoms 8 months after application (Fig. 6, E and G, and fig. S9). The treated trees looked much healthier than the non-treated control despite both were 100% infected (Fig. 6G). Because symptomatic leaves demonstrated significantly more cell death than asymptomatic leaves based on trypan blue staining (Fig. 3a) and TEM observation (fig. S2), we used ratio of symptomatic leaves vs total leaves as an indicator of cell death caused by HLB. GA treatment significantly reduced the percentage of symptomatic leaves (Fig. 6, E and G, and fig. S9), indicating reduced cell death of sieve element and companion cells in treated leaves. In addition, foliar spray of GA on HLB positive 6-year-old *C. sinensis* var. ‘Valencia’ and var. ‘Vernia’ significantly promoted plant growth including tree height, trunk diameter and canopy volume at eight months after application (fig. S10). It is probable that GA reduces HLB symptoms via its direct effect on both mitigating ROS (Fig. 6A) and promoting plant growth (fig. S10).

**Fig. 6.**
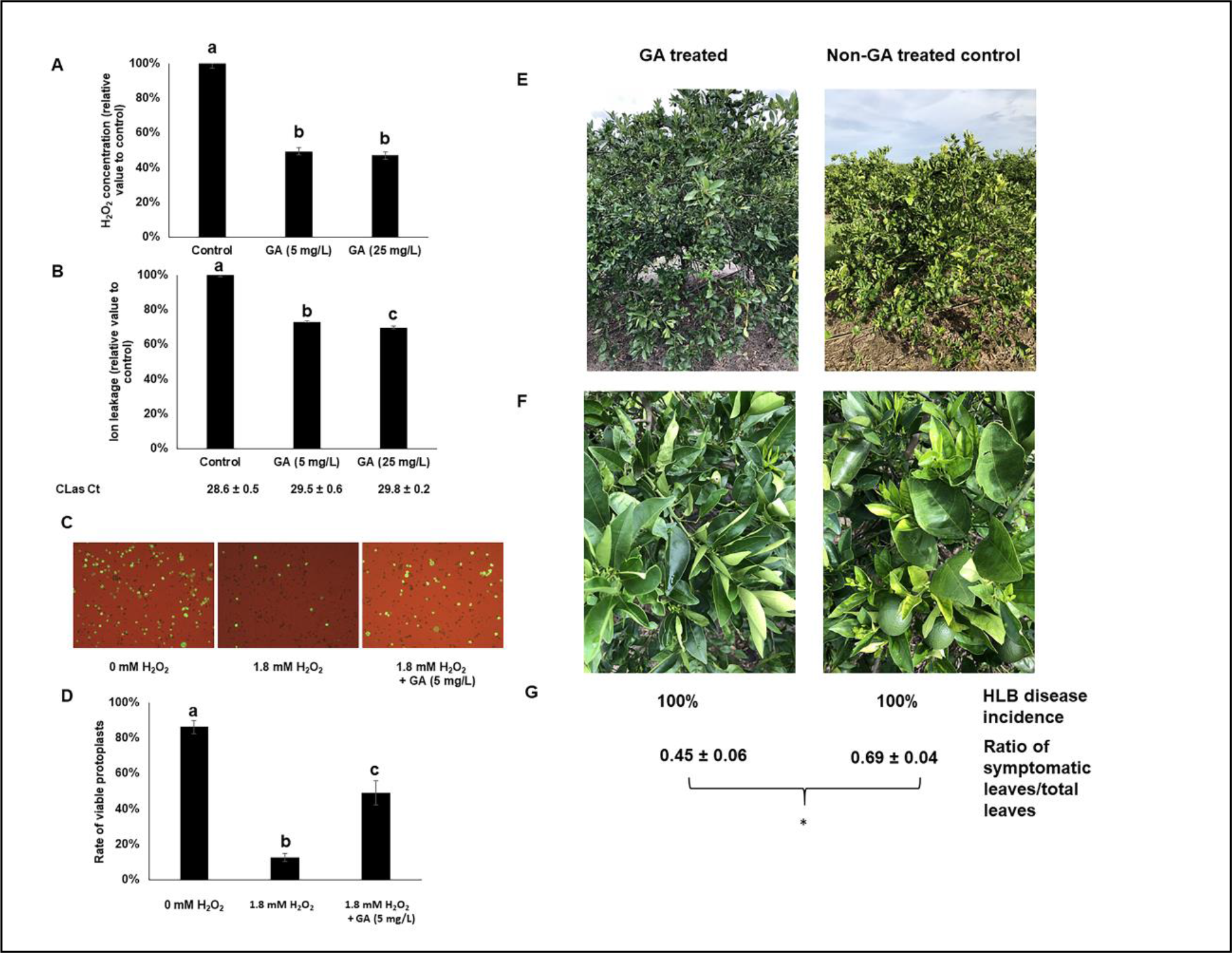
Immunoregulator gibberellin (GA) suppresses HLB development. (A and B) GA suppresses ROS mediated cell death. HLB-positive *C. sinensis* trees were treated with GA via foliar spray weekly for six weeks. The exudates extracted from phloem enriched bark tissues were used for detection of H2O2 (A) and ion leakage (B). n=4. Ct values of CLas of the tested samples were indicated. (C and D) GA suppresses cell death of *C. sinensis* protoplast cells. Freshly prepared protoplast cells of *C. sinensis* were treated with H2O2 with or without GA (5 mg/L) for 24 h and tested for viability via fluorescein diacetate (FDA) staining. Statistical differences (A, B, and D) were analyzed using one way ANOVA with Bonferroni Correction (*P*<0.05). Different letters above the columns indicate statistical differences (*P*<0.05). (E-G) Foliar spray of GA suppresses HLB symptoms. *C. sinensis* ‘Valencia’ blocks were treated with GA (1,247 ppm) in November 2020. Nearby blocks of *C. sinensis* ‘Valencia’ that were not treated with GA were used as negative controls. Symptoms, HLB disease incidence and ratio of symptomatic leaves/total leaves were investigated in June 2021. (E) Representative whole trees. (F) Representative sections. (G) HLB disease incidence and ratio of symptomatic leaves vs total leaves in different treatments. Pictures were taken at the same day in June 2021. * indicates *P* value < 0.05 based on Student’s t-test.

To determine whether mitigating ROS can directly halt or reduce HLB symptoms, we conducted weekly foliar spray of antioxidants uric acid (1.8 mM) and rutin (0.6 mM) on HLB positive *C. sinensis*. Remarkably, 6weeks after the first treatment, both antioxidants significantly reduced HLB symptoms compared to that before treatment, whereas the plants treated with water became more symptomatic in the same duration (fig. S8). Taken together, our data suggest that suppression of ROS mediated cell death mitigates HLB symptoms. Consequently, we have established the causal relationship that CLas triggers ROS production in the phloem tissues, which subsequently causes cell death of phloem tissues, leading to HLB symptoms.

## Discussion

In this study, we demonstrate that citrus HLB is an immune-mediated disease. CLas induces a systemic chronic immune response, mimicking systemic chronic inflammation diseases of human ^35^. Systemic chronic inflammation diseases have been suggested to result from collateral damage to tissues and organs over time by oxidative stress ^36^. ROS concentrations triggered by CLas infection are above the threshold needed to induce cell death. Persistent induction of ROS by systemic CLas infection leads to systemic cell death of phloem tissues and other effects owing to diverse roles of ROS, which subsequently affect phloem function, hormone synthesis and transportation, metabolic transportation, and rerouting energy to immune response rather than to growth. This hypothesis can explain most HLB phenomena. For instance, phloem dysfunction resulting from death of companion and sieve element cells may lead to starch accumulation, and blotchy mottle symptoms. Hardened leaves perhaps result from the action of ROS since ROS are known to directly cause strengthening of host cell walls ^25^. Both cell death of the phloem tissues and reduced transportation of photosynthates may be responsible for root decay. Stunt growth probably results from the direct effect of ROS, reduced transportation of carbohydrates and hormones. The detailed molecular mechanism of how CLas activates immune response remains unknown. We anticipate that cytoplasmic receptors, such as nucleotide-binding leucine-rich repeat (NLR) proteins are mainly responsible for intracellular detection of CLas through recognition of PAMPs inside companion and sieve element cells. It is probable that immune-mediated diseases, even though which have not been previously known for plants, are prevalent for the Plantae Kingdom, such as diseases caused by phloem-colonizing pathogens including bacteria (e.g., *Ca*. Liberibacter, *Spiroplasma*, and *Ca*. Phytoplasma), viruses, and fungi and some non-phloem colonizing pathogens.

Knowing citrus HLB is an immune mediated disease helps guide the battle against this notorious disease. It is projected that horticultural approaches that suppress oxidative stress can provide immediate help to alleviate the immune-mediated damages caused by CLas in HLB endemic citrus production areas. These approaches include optimized usage of plant growth hormones, such as GA and brassinosteroids ^37^, and optimized antioxidants treatment. Even though we did not test the effect of nutritional modulation of immune function on HLB in this study, citrus growers in Florida have observed that modulation of macronutrients (N, P, and K) and micronutrients (e.g., B, Cu, Fe, Mn, Mo, Ni, Se and Zn) reduces HLB symptoms. This is consistent with that a deficiency of the macronutrients leads to oxidative stress ^38^, whereas micronutrients (B, Cu, Fe, Mn, Mo, Ni, Se and Zn) at low concentrations activate antioxidative enzymes ^39^. Antioxidants (e.g., uric acid), growth hormones (e.g., GA) and nutritional modulation (e.g., micronutrients) directly alleviate the oxidative stress to reduce cell death of the phloem tissues to mitigate HLB symptoms. Moreover, growth hormones and micronutrients promote new growth, which decreases the ratio of dead cells in phloem tissues, further mitigating HLB symptoms. The horticultural measures used to mitigate ROS and cell death are expensive and unable to reduce or eliminate CLas inoculum, thus are not recommended for citrus production areas with low HLB incidence. For those areas, region-wide comprehensive implementation of roguing infected trees, tree replacement, and insecticide applications has been shown to successfully control citrus HLB ^6, 40^. Genetic improvements that enhance plant tolerance of oxidative stress, prevent overproduction of ROS, or evade recognition of CLas may generate HLB resistant/tolerant citrus varieties. In particular, we can conduct companion cell- or phloem-specific overexpression of antioxidant enzymes (such as superoxide dismutase, catalases, glutathione peroxidases, ascorbate peroxidase, and glutathione reductase) using CRISPR gene editing, transgenic, or cisgenic approaches or citrus tristeza virus vectors). Additionally, gene editing of the NLRs responsible for recognizing CLas might also render citrus tolerant to HLB. It is plausible that some of the HLB tolerant citrus genotypes might not recognize PAMPs or proteins of CLas or the recognition is too weak to reach the ROS threshold to cause cell death, thus avoiding the immune-mediated damages. On the other hand, some HLB-tolerant citrus genotypes might be superior in antioxidant mechanism or phloem regeneration. Intriguingly, it was reported that HLB-tolerant varieties contain higher levels of antioxidant capacities than the susceptible varieties ^41^. ‘Sugar Belle’ mandarin, which is HLB-tolerant, has more vigorous phloem regeneration than the HLB-susceptible *C. sinensis* varieties^42^. In summary, citrus HLB is an immune-mediated disease and mitigating ROS via antioxidant mechanisms and promoting new growth both can reduce cell death of the phloem tissues, thus controlling HLB.

## Acknowledgements

The research has been supported by Florida Citrus Initiative, Florida Citrus Research and Development Foundation, USDA National Institute of Food and Agriculture grant # 2018-70016-27412, #2016-70016-24833, and #2019-70016-29796.

## Author Contributions

N.W. conceptualized, designed the experiments, and supervised the project. WXM performed ROS related experiments; ZQP conducted ion leakage experiments; WXM and ZQP conducted trypan blue staining together; XEH conducted research related to protoplast; SSP tested H2O2 concentrations, callose deposition, and gene expression of young leaves; JYL conducted foliar spray and trunk injection; JX conducted bioinformatic analysis of transcriptomic data; DA conducted TEM analyses; FV conducted callose deposition analyses of mature leaves; ZQP, FV, and YXH conducted overexpression of CLas genes in citrus, tobacco and Arabidopsis; CH was involved in gene expression using RT-qPCR; WTW, YXH, and JYL participated in ROS and ion leakage experiments; DHL participated in field trials. N.W. wrote the manuscript with input from all co-authors.

## Author Information

The authors declare no competing financial interests. Readers are welcome to comment on the online version of the paper. Correspondence and requests for materials should be addressed to N.W.

## Materials and Methods

### Transgenic expression analysis of CLas proteins containing Sec secretion signals and other putative virulence factors

Transgenic expression of CLas genes was conducted as described previously ^1^. For the citrus transformation, CLas genes were amplified without signal peptide sequence and cloned into the binary vector RCsVMV-erGFP-pCAMBIA-1380N-35S-BXKES-3xHA, which has a C-terminal 3×HA tag, to generate the CLas gene overexpression vectors. The resulting binary vectors were transferred into *Agrobacterium tumefaciens* strain EHA105 for citrus transformation. The empty vector (EV) was used in citrus transformation as a negative control. *Agrobacterium*-mediated transformation of epicotyl segments of Duncan grapefruit (*Citrus paradisi*) was carried out as described previously ^2^. Transgenic lines showing kanamycin-resistance and erGFP-specific fluorescence were selected and then micro-grafted in vitro onto one-month-old Carrizo citrange rootstock seedlings. After one month of growth in vitro, the grafted shoots were potted into a peat-based commercial potting medium and acclimated under greenhouse conditions for the phenotype evaluation. Transgenic plants were confirmed by PCR, qRT-PCR at the RNA level, or western blot using HA Tag Antibodies (Sigma-Aldrich, St. Louis, MO).

For the tobacco transformation, *Agrobacterium*-mediated transformation of leaf discs of *Nicotiana tabacum* was carried out to generate the transgenic tobacco ^3^. *A. tumefaciens* strain EHA105 containing the vectors was used for transformation. Transgenic positive shoots showing kanamycin-resistance and erGFP-specific fluorescence were selected and transferred to the rooting medium. Evaluation of the transgenic *N. tabacum* was conducted in a growth chamber. Transgenic plants were confirmed by PCR, qRT-PCR at the RNA level, or western blot using HA Tag Antibodies (Sigma-Aldrich).

For the gene overexpression in *Arabidopsis thaliana.* CLas genes without signal peptides were PCR amplified and cloned into the binary vector pCambia1380-35S-EYFP, which has a C-terminal EYFP protein tag, and transferred into *A. tumefaciens* strain GV2260. *Agrobacterium*-mediated floral dip method was used for the *Arabidopsis* transformation as reported previously^4^. The T1 generation transgenic plants were screened on the Hygromycin B selection medium. Positive plants were further confirmed by PCR and western blot using GFP antibodies (Sigma-Aldrich). The positive T2 generation transgenic plants were evaluated in a growth chamber.

### Plant materials used for investigation of the relationship between CLas infection, immune response, phloem blockage, cell death and HLB symptom development

We used two-year-old CLas-infected and healthy Valencia sweet orange (*Citrus sinensis*) plants maintained in a greenhouse (28°C ± 2°C, relative humidity of 50% ± 5%, natural light period). We selected young flushes from the twigs of HLB-positive sweet orange trees. Both ‘Valencia’ and ‘Hamlin’ sweet orange are *C. sinensis* and susceptible to HLB without observable differences in symptoms. The plant was infected with CLas by graft inoculation and maintained in a greenhouse. The sweet orange trees from the groves were naturally infected by CLas. Healthy plants were maintained in a glasshouse with natural light and without temperature control.

### Quantification of H2O2 concentrations

H2O2 concentrations were quantified following the procedure described elsewhere ^5^. Briefly, leaf samples (0.5 g) were grinded in 0.1% (w/v) trichloroacetic acid (TCA) and centrifuged at 12,000 g for 15 min at 4°C. The supernatant (0.3 ml) was mixed with 1.7 ml 1.0 M potassium phosphate buffer (pH 7.0) and 1.0 ml of 1.0 M potassium iodide solution, then incubated for 5 min before measuring the absorbance of the oxidation product at 390 nm. H2O2 concentrations were calculated using a standard curve prepared with known concentrations of H2O2 and expressed in µmol/g fresh weight. For measuring H2O2 concentrations in the exudates of phloem enriched bark tissues, the same procedure was used except the TCA step and H2O2 concentrations were expressed in mmol/L.

### Ion leakage

Conductivity of the exudates extracted from phloem enriched bark tissues was measured using a CON 700 conductivity/°C/°F bench meter (OAKTON Instruments, Vernon Hills, IL, USA).

### Callose deposition assay

Leaf samples were fixed with FAA (37% formaldehyde/glacial acetic acid/95% ethanol/deionized water at a volume ratio of 50:5:10:35) solution overnight. Samples were embedded in the Tissue Plus O.C.T compound (Thermo-Fisher, Waltham, MA, USA), sectioned with a Harris Cryostat Microtome (International Equipment, Boston, MA, USA) and stained with 0.005% aniline blue solution prior to analysis. Samples were observed in an Olympus BX61 epifluorescence microscope (Olympus Corporation, Center Valley, PA, USA). Callose spots were counted per slide area for all sample types.

### CLas quantification using qPCR

Tissues (100 mg) were homogenized into powders using a TissueLyser II (Qiagen, Valencia, CA, USA). DNA was extracted using the DNeasy Plant kit (Qiagen), following the manufacturer’s instructions, and eluted in 100 µL nuclease free water. DNA concentration was measured using a Synergy LX plate reader (BioTek, Winooski, VT, USA). Quantification of CLas in plant tissues was performed as described elsewhere ^6^. Briefly, qPCR was carried out with primers and probe for CLas ^7^. qPCR assays were performed with QuantiStudio3 (Thermo Fisher, Waltham, MA) using the Quantitec Probe PCR Master Mix (Qiagen) in a 25-µl reaction. The standard amplification protocol was 95°C for 10 min followed by 40 cycles at 95°C for 15 s and 60°C for 60 s. All reactions were conducted in triplicate with CLas positive and water controls. Quantification of CLas was conducted using the equation Y = -0.288 ✕ (CLas Ct) + 11.607 ^8^.

### Starch assay

The samples (100 mg) were powdered using a TissueLyser II (Qiagen, Hilden, Germany). The powdered samples were used to quantify the starch. The starch estimation was performed using the Total Starch Assay Kit (AA/AMG) (Megazyme, Bray, Ireland) as instructed by the manufacturer. The experiments were repeated thrice with similar result.

### Statistical analyses

All statistical analyses were performed using SAS statistical software (Version 9.4, SAS Institute, Cary, NC, USA).

### Gene expression assays using reverse transcription quantitative PCR (RT-qPCR)

Total RNA was extracted using the RNeasy Plant Mini Kit (Qiagen), according to manufacturer’s instructions. cDNA was synthesized with Quantitec Reverse Transcription Kit (Qiagen) according to manufacturer’s instructions and diluted 10 times for RT-qPCR. Reactions were carried out by adding 1 µL of cDNA, 1 µL of each specific primer, 7 µL water and 10 µL Fast SYBR Green Master Mix (Thermo Fisher Scientific, Waltham, MA, USA) performed with QuantiStudio3 (Thermo Fisher) using the standard fast protocol of 95°C for 20 s followed by 40 cycles of 95°C for 1 s and 60°C for 20 s. Denaturation protocol consisted of 95°C for 1 s, 60°C for 20 s and a final dissociation step of 95°C. Relative gene expression was calculated using the method described previously ^9^. CsGAPDH was used as an endogenous control.

### TEM analysis

Small sections of the leaf and stem samples were collected under a stereomicroscope (Swift Table Stereo Zoom Microscope, Carlsbad, CA, USA). The leaf samples were transferred to 3% glutaraldehyde overnight at 4°C for fixation. Then, the samples were postfixed in 2% osmium tetroxide prepared in 3% glutaraldehyde for 4 h at room temperature in a fume hood. The samples were dehydrated by sequential treatment with 10%, 20%, 30%, 40%, 50%, 60%, 70%, 80%, 90% and 100% (thrice) acetone for 10 min each. The leaf samples were incubated sequentially in 50%, 75% and 100% (twice) Spurr’s low-viscosity epoxy resin prepared in acetone for 8 h each. One-micrometer sections were cut with glass knives using an ultramicrotome followed by staining with methylene blue/azure A for 30 sec and basic fuchsin (0.1 g in 10 ml of 50% ethanol) for 30 sec. The sections were observed under a Leitz Laborlux S compound microscope (Leica Microsystems, Wetzlar, Germany) for the right spot with a vascular system. The same blocks were trimmed with a surgical blade and then sectioned to 0.1 µm using a diamond knife under an ultramicrotome. The thin sections were collected on 200-mesh copper grids. The samples were stained with 2% aqueous uranyl acetate for 5 min, washed in water, and again stained with lead citrate followed by water wash. The micrographs were prepared and analyzed using a Morgagni 268 (FEI Company, Hillsboro, OR, USA) transmission electron microscope equipped with an AMT digital camera (Advanced Microscopy Techniques Corp., Danvers, MA, USA).

### Trypan-blue staining

Trypan-blue staining was conducted as described by Fernández-Bautista et al. ^10^.

### Monitoring ROS formation and localization in phloem tissues by use of the fluorescent probe 2’,7’-dichlorodihydrofluorescein diacetate (H2DCFDA) and confocal laser microscopy

Four CLas-infected branches collected from HLB-positive *C. sinensis* ‘Valencia’ trees and four CLas-free branches collected from healthy *C. sinensis* ‘Valencia’ trees were collected and placed in glass test tube with 30 mL water containing 10 μM 2’,7’-dichlorodihydrofluorescein diacetate (H2DCFDA). Four leaves/branch at the top were kept to facilitate transpiration. Glass tubes were wrapped with aluminum foil and kept in room temperature for 24 hours. Bark was peeled from the stem section that was submerged in water and placed on slide with inner side upwards. HLB-positive and healthy branches were also incubated in water without H2DCFDA as controls. 2’,7’-dichlorofluorescein (DCF) fluorescence was visualized by confocal laser scanning microscopy (CLSM) (Leica TCS-SP5, Mannheim, Germany) with excitation/emission at 495 nm/525 nm.

### Trunk injection of HLB-positive 5-year-old *C. sinensis* trees

Trunk injection was conducted as described elsewhere ^11^. For each tree, approximately 0.4 g streptomycin sulfate (laboratory grade; Thermo Fisher Scientific) at 5 g/L was injected into the trunk. The amount of streptomycin injected was calculated to reach the concentration needed to kill CLas in planta based on the canopy volume ^11^.

### Exudates of phloem enriched bark tissues for H2O2 and ion leakage assays

Exudates of phloem enriched bark tissues were extracted from stems following the procedure described elsewhere ^12^. Stems were collected from small branches of spring spouts with mildly symptomatic leaves.

### Protoplast

Protoplast cells of *C. sinensis* ‘Hamlin’ were prepared as described by ^13^. Embryogenic calli were subcultured on solid MT (Murashige and Tucker) media (Phytotech) every 2 weeks. From the maintained calli suspension, cells were prepared and maintained in DOG liquid media as described elsewhere ^14^. The final isolated protoplast cells were suspended in W5 solution (154 mM NaCl,125 mM CaCl2, 5 mM KCl, 2mM MES at pH 5.7) at 1 × 10^7^ cells/ml for different treatments.

### Foliar spray with antioxidants and GA

Five-year-old Valencia sweet orange trees with similar symptoms were used for foliar spray treatments. All trees in the grove were HLB-positive. The experiment was a completely randomized design with 5 treatments. Each treatment consisted of four trees. The treatments were applied by foliar spray with 2.5 mL/L of Induce non-ionic surfactant (Helena Ag, Collier, TN, USA). One liter of solution per plant were applied at approximately 400 kPa using a handheld pump sprayer. This pressure resulted a fine mist and was sufficient to produce runoff from the leaves to ensure complete coverage. Individual treatments were applied to the various trees as follows: uric acid (1.8 mM), rutin (0.6 mM), GA (5 mg/L), and GA (25 mg/L). Water was used as the negative control. Foliar spray was conducted in the evening to facilitate absorption. The chemicals GA, and uric acid were purchased from Fisher Scientific. Rutin was purchased from Sigma-Aldrich (St. Louis, MO, USA).

### GA treatment via foliar spray

GA foliar spay was conducted in the first week of November, 2020. For the GA application, 20 ounces of Pro Gibb LV (Valent U.S.A. LLC, Walnut Creek, CA, USA) was mixed with water in a 100 Gal tank. 64 ounces of WIDESPREAD MAX (A.I. organosilicone) was included as the surfactant for leaf spray with airblast. Applications were conducted during night. One block of Valencia sweet orange on rootstock 942 was treated with GA, whereas the nearby Valencia/942 block was not treated with GA as a negative control. In addition, one block of Vernia sweet orange on X639 rootstock was treated with GA with one nearby Vernia/X639 block as a negative control. All blocks are 10 acres or more.

### Evaluation of citrus tree growth and HLB symptoms in response to GA treatment

Tree growth was evaluated by estimating trunk diameter, tree height (TH), and canopy volume (CV) on both GA treated and untreated control trees. For each treatment group, a total of 10 trees (n =10) were randomly selected for the evaluation. A digital caliper (Fowler, Newton, MA) was used to take two measurements of trunk diameter (north-south and east-west orientation) at ∼20 cm above the ground. A tape measure was used to measure the TH above the ground from the soil surface to the apical point of the plant. CV was estimated by taking the average of two independent measurements of the diameter of the canopy at different directions (north-south and east-west). The CV was estimated using the equation: V = (2/3) × p × h × (d/2)^2^, where h is the TH and d is the average diameter of the tree canopy ^15^. All statistical analyses were performed using SAS V9.4 (SAS Institute Inc., Cary, NC). The data were first tested for normality and homogeneity of variance using the Shapiro-Wilk’s test and Levene’s test, respectively. A Student’s two-tailed t test was performed to explore differences between GA treated and untreated control trees in growth performance traits.

HLB disease incidence in different treatments was evaluated by randomly checking 200 trees/treatment. Ratio of symptomatic leaves vs total leaves in different treatments were investigated by evaluating 3 groups of branches/treatment with each group containing 16 branches that were selected randomly from 8 trees (2 branches/tree).

### Data analyses of RNA-seq data

To generate the comprehensive expression pattern of sweet orange in response to CLas infection, we collected 15 microarray and 9 RNA-seq data sets from NCBI SRA and GEO databases (Table S2). The differentially expressed genes (DEGs) were determined using Limma ^16^ and DESeq2 ^17^ packages in R for microarray and RNA-seq data, respectively (adjusted p value <0.05 and |log2 fold change| >1). Gene ontology (GO) term enrichment of DEGs was conducted using agriGO v2.0: a GO analysis toolkit for the agricultural community ^18^ using the singular enrichment analysis tool. The heatmap plots were drawn using the gplots package in R program ^19^

---

**Fig. S1.**
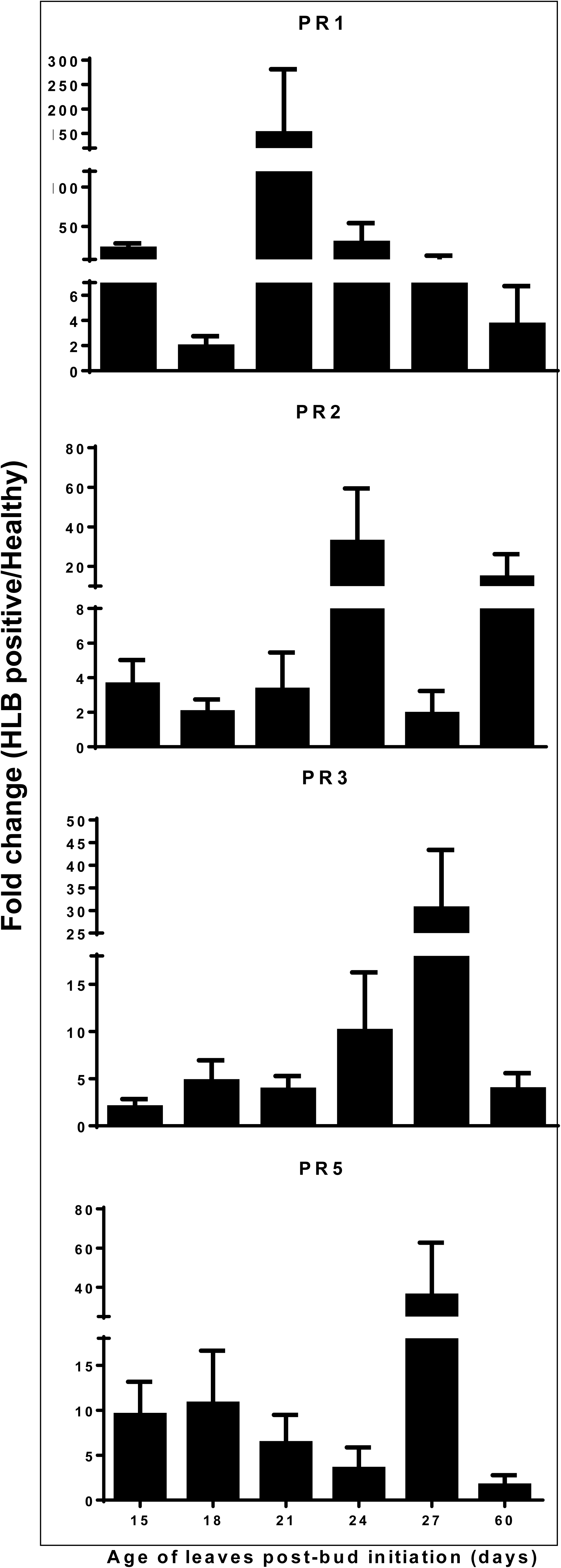
Temporal expression of immune-related genes in response to CLas infection of young flush of *C. sinensis*. The reverse transcription-quantitative PCR (RT-qPCR) analysis was conducted using young leaves from three two-year-old HLB positive Valencia sweet orange trees compared with that of three healthy Valencia trees with one tree as one biological replicate.

**Fig. S2.**
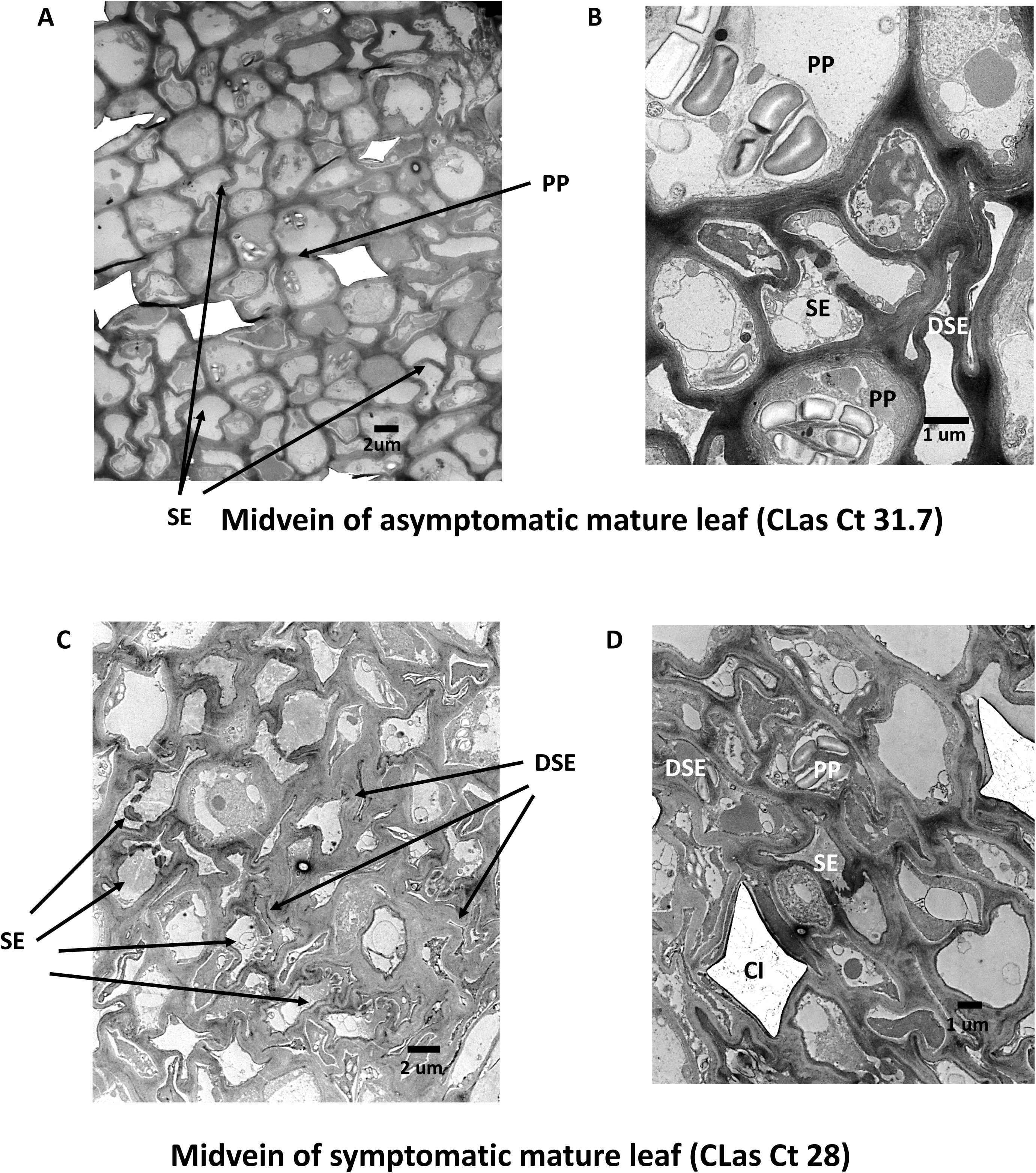
TEM observation of the midribs of mature leaves of *C. sinensis* trees grown in the field that are HLB positive. (A and B) Midvein of asymptomatic mature leaf. (C and D) Midvein of symptomatic mature leaf. CLas titers were indicated by Ct values for the samples used. Scale bar for each picture is included. SE: sieve element. DSE: dead sieve element cells. PP: parenchyma cells. CI: calcium oxalate crystal idioblas.

**Fig. S3.**
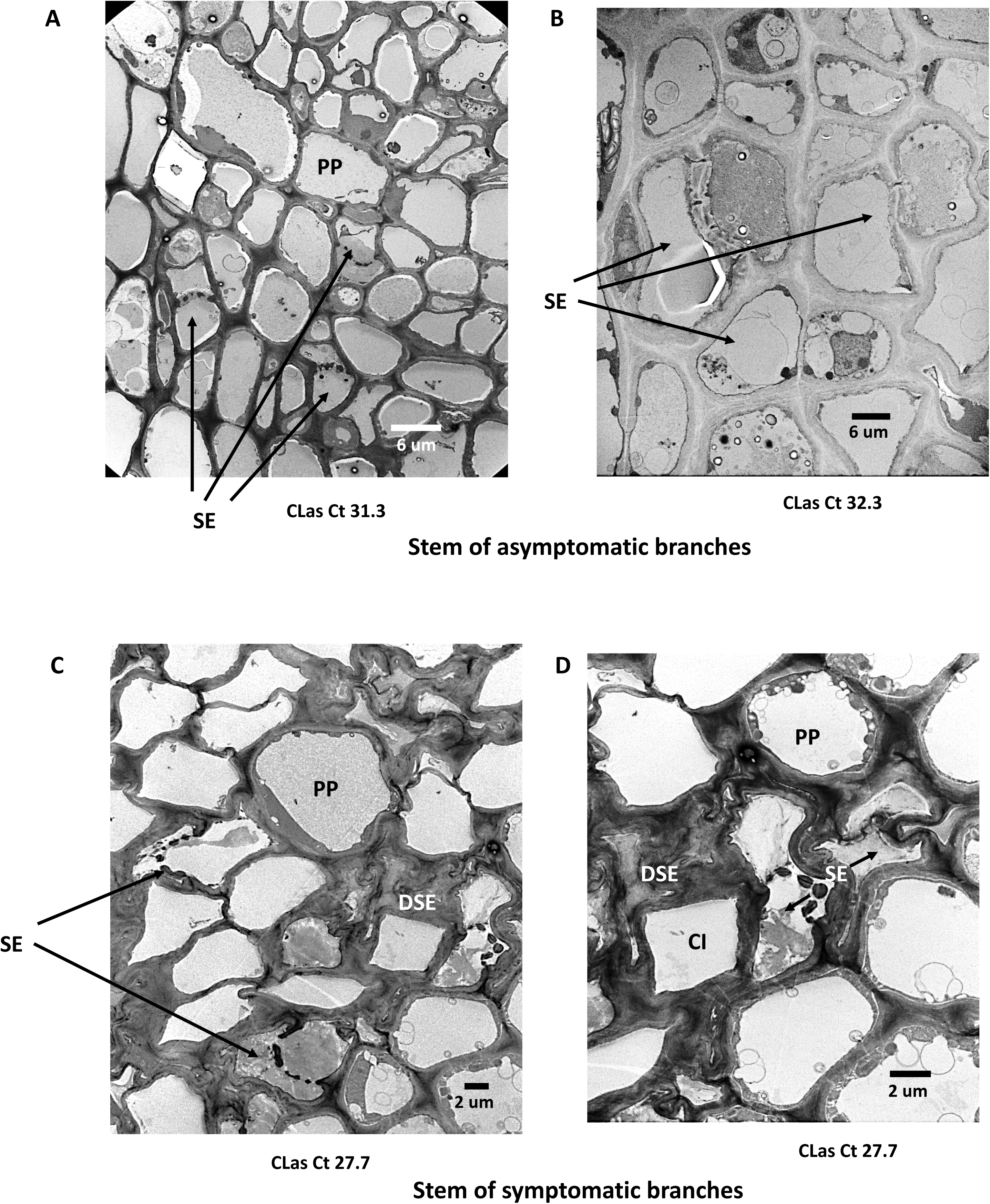
TEM observation of the stem tissues of HLB positive *C. sinensis* trees grown in the field. Stems were collected from branches without HLB symptoms (A and B) and branches with HLB symptoms (C and D). Scale bar for each picture is included. SE: sieve element. DSE: dead sieve element cells. PP: parenchyma cells. CI: calcium oxalate crystal idioblas. CLas titers in the tested samples were indicated by Ct values.

**Fig. S4.**
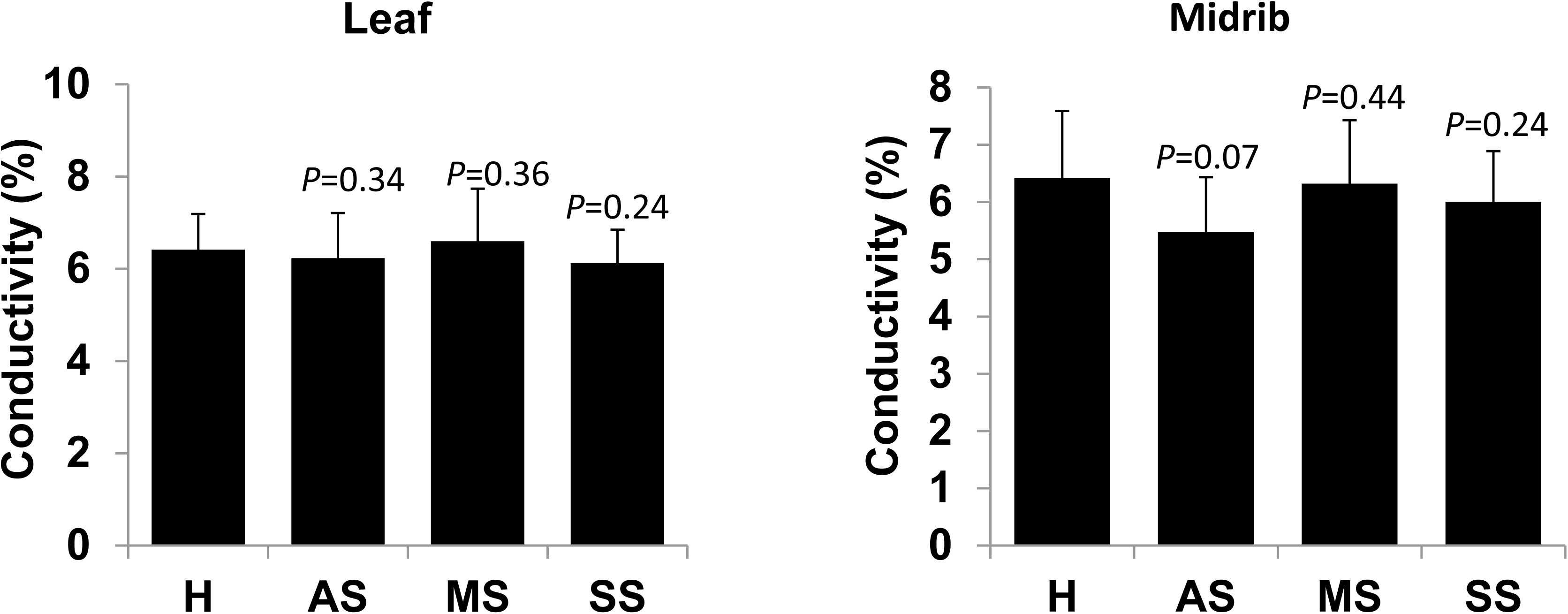
Comparison of ion leakage activities of leaves of CLas-negative and CLas-infected sweet orange trees. H: healthy leaves. AS: asymptomatic leaves. MS: leaves with mild symptoms. SS: leaves with severe symptoms. Healthy leaves were collected from CLas-free sweet orange plants. AS, MS, and SS were collected from CLas positives sweet orange trees in the groves. For statistical significance tests, Student’s t-test was conducted (n=9). Mean and standard deviation are shown. P values for comparisons against CLas negative samples were shown above each column.

**Fig. S5.**
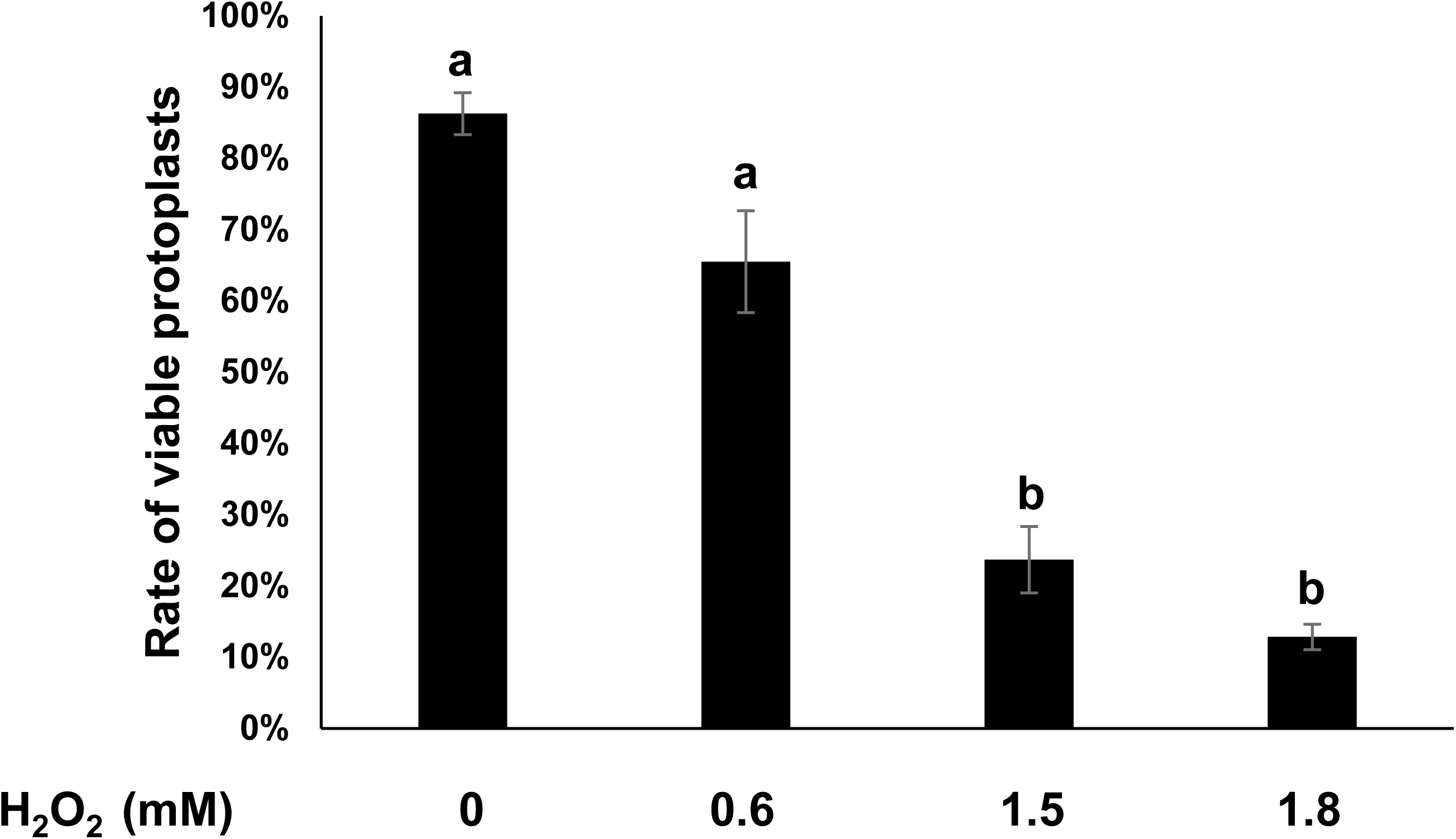
H2O2 kills protoplast cells of *C. sinensis*. Freshly prepared protoplast cells of *C. sinensis* were treated with different concentrations of H2O2 for 24 h and tested for viability via fluorescein diacetate (FDA) staining. Each experiment contains three biological replicates. Mean and SD were shown. Statistical differences were analyzed using one way ANOVA with Bonferroni Correction (*P*<0.05). Different letters above the columns indicate statistical differences (*P*<0.05).

**Fig. S6.**
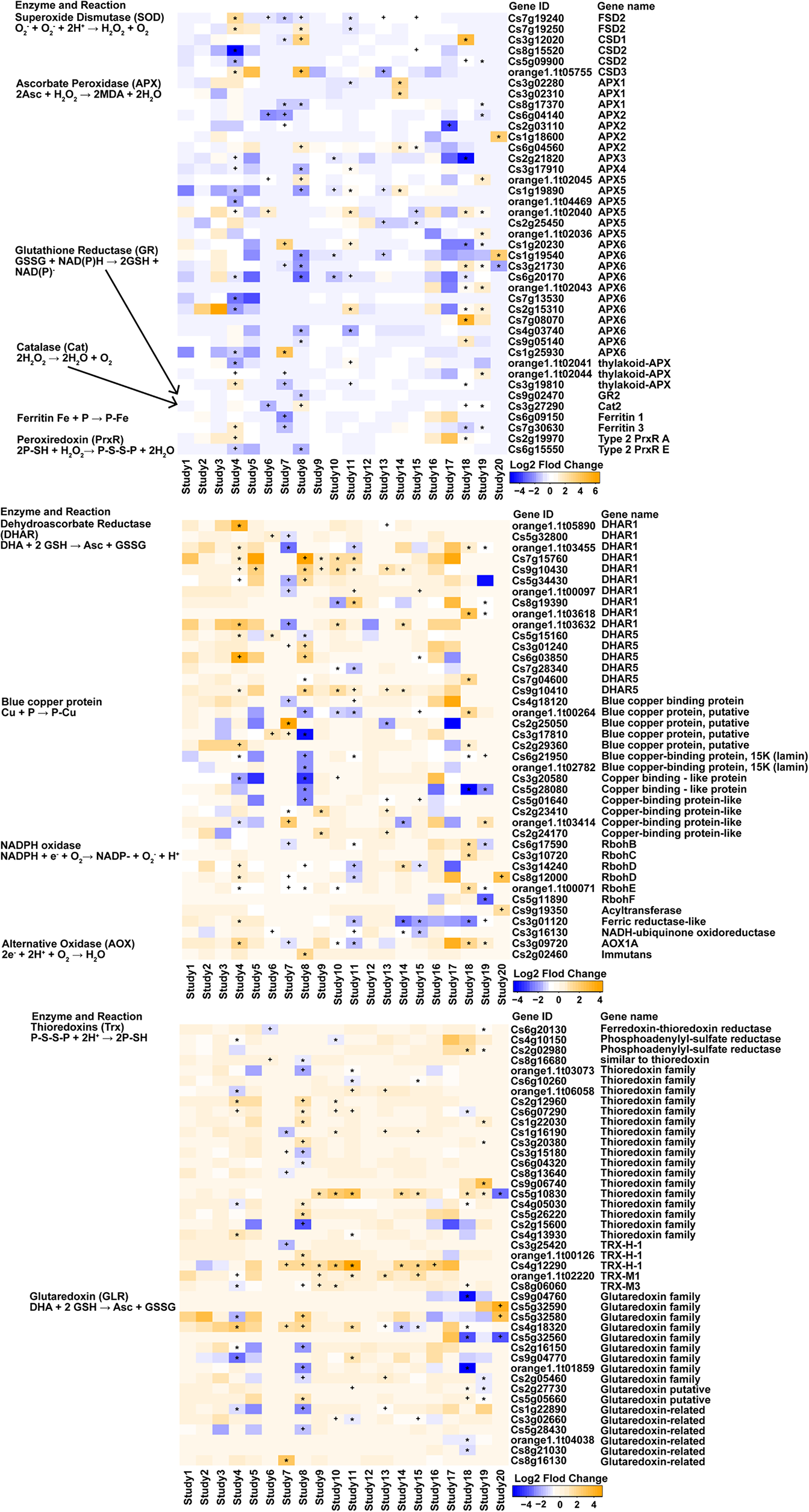
The expression profile of ROS-related genes in response to CLas infection in 9 previous studies. Affiliation of each gene is indicated in *brackets*. *Orange* denotes “higher in CLas infected than CLas negative samples” while blue denotes “higher in CLas negative than CLas infected samples”. The asterisk denotes *P* value < 0.01; the plus sign denotes *P* value < 0.05. The information about the 9 previous studies is listed in Table S2.

**Fig. S7.**
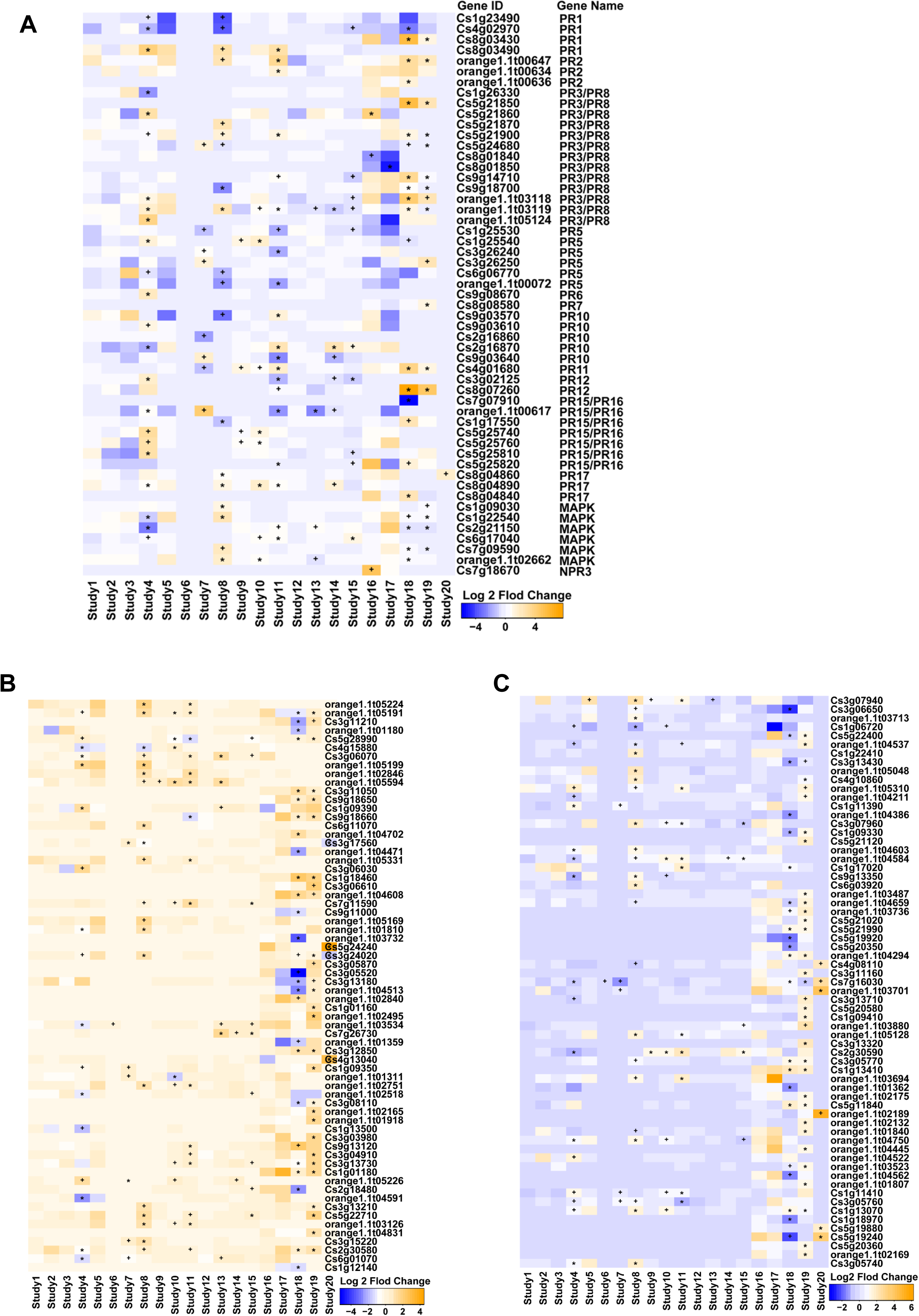
The expression profile of immune related genes, including PR, MAPK and NBS-LRR genes in response to CLas infection in 9 previous studies. Affiliation of each gene is indicated in *brackets*. *Orange* denotes “higher in CLas infected than CLas negative samples” while blue denotes “higher in CLas negative than CLas infected samples”. The asterisk denotes *P* value < 0.01; the plus sign denotes *P* value < 0.05. (A) PR genes, and MAPK genes. (B and C) NBS-LRR genes. The information about the 9 previous studies is listed in Table S2.

**Fig. S8.**
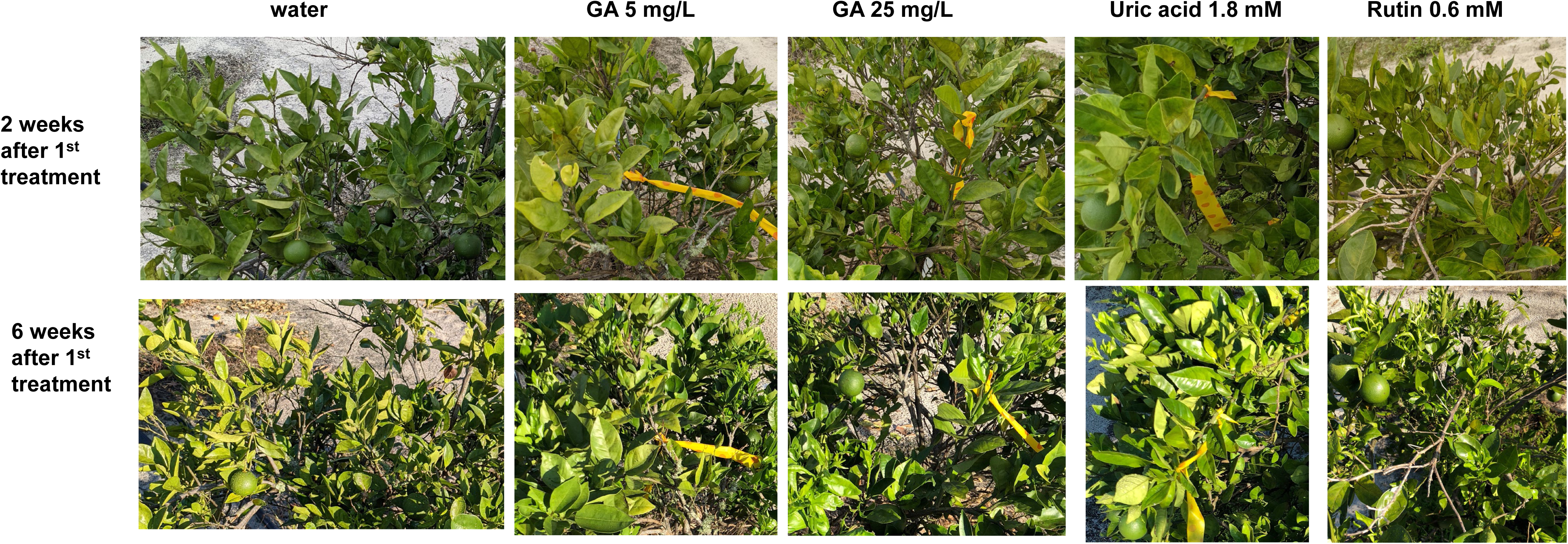
Effect of GA and antioxidants treatment on HLB symptoms. To test the effect of GA treatment on HLB symptoms, HLB positive *C. sinensis* ‘Valencia’ trees were treated with GA and antioxidants (uric acid and rutin) via foliar spray weekly. Representative branches were selected to demonstrate symptom changes.

**Fig. S9.**
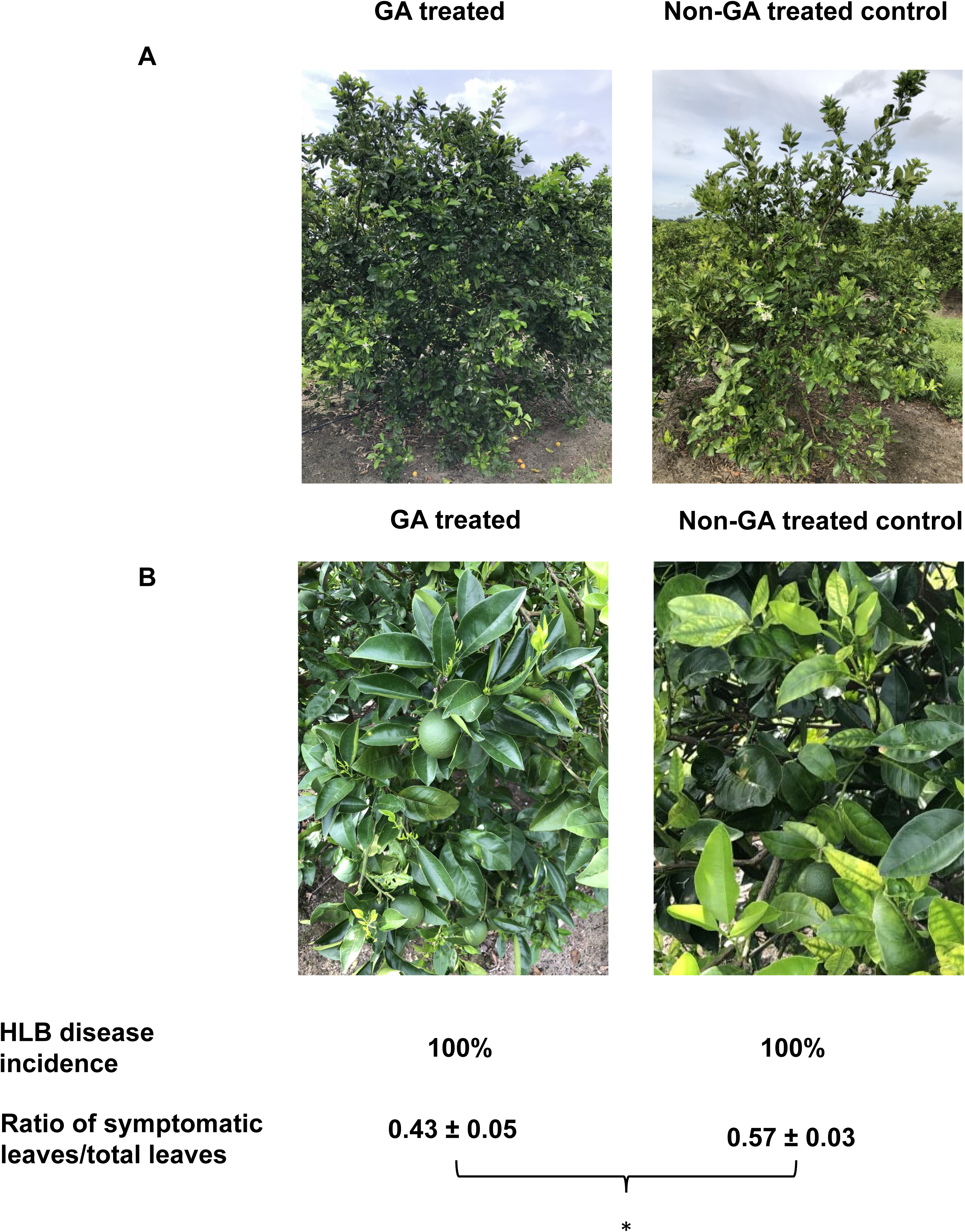
Gibberellin (GA) treatment of *C. sinensis* suppresses HLB. *C. sinensis* ‘Vernia’ blocks were treated with GA (1247 ppm) in November 2020. Nearby blocks of *C. sinensis* ‘Vernia’ that were not treated with GA were used as negative controls. Symptoms, HLB disease incidence and ratio of symptomatic leaves/total leaves were investigated in June 2021. (A) Representative whole trees. (B) Representative sections. (C) HLB disease incidence and ratio of symptomatic leaves vs total leaves in different treatments. Pictures were taken at the same day in June 2021. * indicates *P* value < 0.05 based on Student’s t-test.

**Fig. S10.**
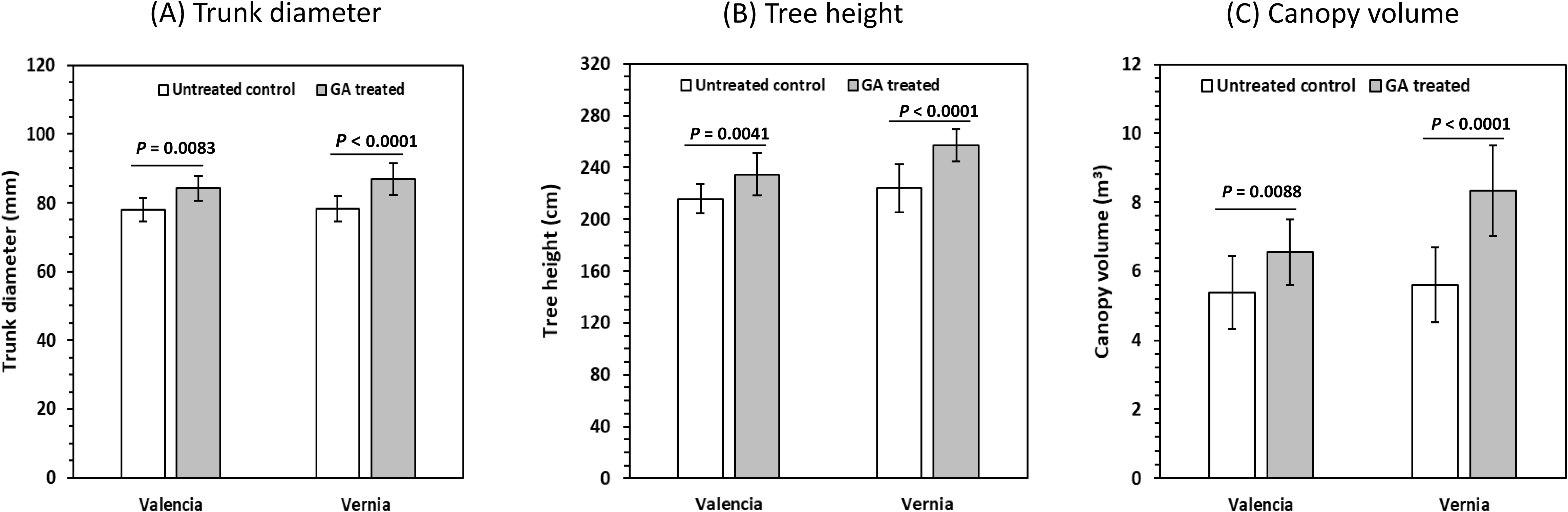
Growth performance of citrus trees (two cultivars: Valencia and Vernia) treated by GA in commercial groves in FL. *C. sinensis* blocks were treated with GA (1247 ppm) in November 2020. Nearby blocks of *C. sinensis* that were not treated with GA were used as negative controls. Trees were investigated in June 2021. (A) Trunk diameter measured at ∼20 cm above the ground. (B) Tree height above the ground from the soil surface to the apical point of the plant. (C) Canopy volume of the GA treated and untreated control trees. Data shown are means of 10 replicated trees (n = 10). The error bars are standard deviations. P values of Student’s two-tailed t tests were presented and a significant difference between GA treated, and untreated control trees was determined at *P* < 0.05.

**Table S1:** Overexpression of CLas proteins containing Sec-section signals and other predicated virulence factors in *Arabidopsis*, *Citrus* and *Nicotiana*.

| Locus | Annotation | SDE | Overexpressed in *Arabidopsis thaliana* |  |  | Overexpressed in *Citrus* |  |  | Overexpressed in *N. tabacum* |  |  | Reference |
| --- | --- | --- | --- | --- | --- | --- | --- | --- | --- | --- | --- | --- |
|  |  |  | PCR | WB | Phenotype | PCR | WB | Phenotype | PCR | WB | Phenotype |  |
| CLIBASIA_00460 |  | Y | + | + | Normal |  |  |  |  |  |  |  |
| CLIBASIA_00530 |  | Y | + | + | Normal |  |  |  |  |  |  |  |
| CLIBASIA_02145 |  | Y | + | - | Normal |  |  |  |  |  |  |  |
| CLIBASIA_02215 |  | Y | + | + | Normal |  |  |  |  |  |  |  |
| CLIBASIA_02470 |  | Y | + | - | Normal |  |  |  |  |  |  |  |
| CLIBASIA_02935 |  | Y | + | - | Normal |  |  |  | + | - | Normal |  |
| CLIBASIA_03695 |  | Y | + | + | Normal |  |  |  |  |  |  |  |
| CLIBASIA_03975 |  | Y | + | - | Normal |  |  |  |  |  |  |  |
| CLIBASIA_04320 |  | Y | + | - | Normal |  |  |  |  |  |  |  |
| CLIBASIA_04580 |  | Y | + | - | Normal |  |  |  |  |  |  |  |
| CLIBASIA_04735 |  | Y | + | - | Normal |  |  |  |  |  |  |  |
| CLIBASIA_05115 | predicated virulence factor | Y | + | + | Normal |  |  |  |  |  |  |  |
| CLIBASIA_05320 |  | Y | + | - | Normal |  |  |  |  |  |  |  |
| CLIBASIA_05330 |  | Y | + | + | Normal |  |  |  |  |  |  |  |
| CLIBASIA_04260 | OmpA |  | + | + | Normal |  |  |  |  |  |  |  |
| CLIBASIA_03315 | predicated virulence factor |  | + | + | Normal |  |  |  |  |  |  |  |
| CLIBASIA_03105 | Flp/Fap pilin component |  | + | + | Normal |  |  |  |  |  |  |  |
| CLIBASIA_05640 |  | Y | + | + | Normal |  |  |  |  |  |  |  |
| CLIBASIA_02305 |  | Y | + | + | Normal |  |  |  |  |  |  |  |
| CLIBASIA_03085 |  | Y | + | + | Normal |  |  |  |  |  |  |  |
| CLIBASIA_00420 | outer membrane lipoprotein |  | + | + | Normal |  |  |  |  |  |  |  |
| CLIBASIA_01300 | predicated virulence factor |  | + | - | Normal |  |  |  |  |  |  |  |
| CLIBASIA_01345 | serralysin |  | + | - | Normal |  |  |  | + | - | Normal |  |
| CLIBASIA_02425 | outer membrane protein |  | + | - | Normal |  |  |  |  |  |  |  |
| CLIBASIA_03295 | predicated virulence factor |  | + | - | Normal |  |  |  |  |  |  |  |
| CLIBASIA_04055 | predicated virulence factor |  | + | + | Normal |  |  |  |  |  |  |  |
| CLIBASIA_04520 | predicated virulence factor |  | + | - | Normal |  |  |  | + | - | Normal |  |
| CLIBASIA_04410 |  | Y | + | + | Normal |  |  |  |  |  |  |  |
| CLIBASIA_05150 |  | Y | + | + | Normal |  |  |  |  |  |  |  |
| CLIBASIA_01640 | predicated virulence factor |  | + | + | Normal |  |  |  |  |  |  |  |
| CLIBASIA_04865 | predicated virulence factor |  | + | + | Normal |  |  |  |  |  |  |  |
| CLIBASIA_04405 |  | Y | + | + | Normal | + | + | Normal | + | + | Normal |  |
| CLIBASIA_05160 | predicated virulence factor |  | + | + | Normal |  |  |  |  |  |  |  |
| CLIBASIA_05475 | predicated virulence factor |  | + | + | Normal |  |  |  |  |  |  |  |
| CLIBASIA_04025 | SDE15 | Y | + | + | Normal | + | + | Normal | + | + | Normal | 1 |
| CLIBASIA_05315 | SDE1 | Y |  |  |  | + | + | Normal | + | + | Normal | 20 |
| CLIBASIA_02845 |  | Y |  |  |  | + | + | Normal | + | + | Normal |  |
| CLIBASIA_00255 | SahA |  |  |  |  |  |  |  | + | + | Normal | 21 |
| CLIBASIA_00830 | cytochrome-c oxidase |  |  |  |  |  |  |  | + | - | Normal |  |
| CLIBASIA_04330 |  | Y |  |  |  |  |  |  | + | + | Normal |  |
| CLIBASIA_00470 |  | Y |  |  |  |  |  |  | + | + | Normal |  |
| CLIBASIA_02160 | metalloprotease |  |  |  |  |  |  |  | + | + | Normal |  |
| CLIBASIA_04030 |  | Y |  |  |  |  |  |  | + | + | Normal |  |
| CLIBASIA_00520 |  | Y |  |  |  |  |  |  | + | - | Normal |  |
| CLIBASIA_02395 | TPR repeat-containing protein |  |  |  |  |  |  |  | + | - | Normal |  |
| CLIBASIA_04040 |  | Y |  |  |  |  |  |  | + | - | Normal |  |
| CLIBASIA_01555 | hemolysin |  |  |  |  |  |  |  | + | + | Normal |  |

Note: Normal indicates that the phenotypes of transgenic plants are same as wild type plants. WB: Western blot. SDE: Sec-dependent effector. Y: indicates presence of Sec-secretion signal. “+” indicates presence. “-” indicates absence.

**Table S2:** Transcriptomic studies of sweet orange in response to CLas infection that were used for GO enrichment analysis in this study.

| Study | Genotype | Number of samples | Tissue | Time after CLas infection | GSE Dataset | Sequencing method | Analysis method | Analysis statistic | Condition | Reference |
| --- | --- | --- | --- | --- | --- | --- | --- | --- | --- | --- |
| 1 | Valencia | 6 (3 healthy + 3 HLB) | Stem | 16 months | GSE33004 | Microarray | Limma R Package | Unadjusted $P < 0.05$<br>FC > 2 | greenhouse | Aritua et al., 2013 <sup>22</sup> |
| 2 | Valencia | 5 (2 healthy + 3 HLB) | Leaf | 5-9 weeks | GSE33459 | Microarray | Limma R Package | Unadjusted $P < 0.05$<br>FC > 2 | greenhouse | Albrecht et al., 2008 <sup>23</sup> |
| 3 | Valencia | 5 (2 healthy + 3 HLB) | Leaf | 13-17 weeks | GSE33459 | Microarray | Limma R Package | Unadjusted $P < 0.05$<br>FC > 2 | greenhouse | Albrecht et al., 2008 <sup>23</sup> |
| 4 | Sweet orange | 6 (3 healthy + 3 HLB) | Leaf | 8 months | GSE33003 | Microarray | Limma R Package | Unadjusted $P < 0.05$<br>FC > 2 | greenhouse | Kim et al., 2009 <sup>24</sup> |
| 5 | Madam Vinous | 6 (3 healthy + 3 HLB) | Leaf | 7 months | GSE29633 | Microarray | Limma R Package | Unadjusted $P < 0.05$<br>FC > 2 | greenhouse | Fan et al., 2011 <sup>25</sup> |
| 6 | Madam Vinous | 6 (3 healthy + 3 HLB) | Leaf | 5 weeks | Not available | Microarray | Limma R Package | Unadjusted $P < 0.05$<br>FC > 2 | greenhouse | Fan et al., 2012 <sup>26</sup> |
| 7 | Madam Vinous | 6 (3 healthy + 3 HLB) | Leaf | 17 weeks | Not available | Microarray | Limma R Package | Unadjusted $P < 0.05$<br>FC > 2 | greenhouse | Fan et al., 2012 <sup>26</sup> |
| 8 | Madam Vinous | 6 (3 healthy + 3 HLB) | Leaf | 27 weeks | Not available | Microarray | Limma R Package | Unadjusted $P < 0.05$<br>FC > 2 | greenhouse | Fan et al., 2012 <sup>26</sup> |
| 9 | Hamlin | 8 (4 healthy + 4 HLB) | juice vesicles | — | GSE33373 | Microarray | Limma R Package | Unadjusted $P < 0.05$<br>FC > 2 | Field | Liao et al., 2012 <sup>27</sup> |
| 10 | Hamlin | 8 (4 healthy + 4 HLB) | vascular tissue | — | GSE33373 | Microarray | Limma R Package | Unadjusted $P < 0.05$<br>FC > 2 | Field | Liao et al., 2012 <sup>27</sup> |
| 11 | Hamlin | 8 (4 healthy + 4 HLB) | flavedo | — | GSE33373 | Microarray | Limma R Package | Unadjusted $P < 0.05$<br>FC > 2 | Field | Liao et al., 2012 <sup>27</sup> |
| 12 | Hamlin | 8 (4 healthy + 4 HLB) | Seed | — | GSE33373 | Microarray | Limma R Package | Unadjusted $P < 0.05$<br>FC > 2 | Field | Liao et al., 2012 <sup>27</sup> |
| 13 | Valencia | 8 (4 healthy + 4 HLB) | juice vesicles | — | GSE33373 | Microarray | Limma R Package | Unadjusted $P < 0.05$<br>FC > 2 | Field | Liao et al., 2012 <sup>27</sup> |
| 14 | Valencia | 8 (4 healthy + 4 HLB) | vascular tissue | — | GSE33373 | Microarray | Limma R Package | Unadjusted $P < 0.05$<br>FC > 2 | Field | Liao et al., 2012 <sup>27</sup> |
| 15 | Valencia | 8 (4 healthy + 4 HLB) | flavedo | — | GSE33373 | Microarray | Limma R Package | Unadjusted $P < 0.05$<br>FC > 2 | Field | Liao et al., 2012 <sup>27</sup> |
| 16 | Valencia | 10 (5 healthy + 5 HLB), biologic replicates merged as one sample for each group | Immature fruit | — | SRP022979 | RNA-seq | DESeq | Adjusted $P < 0.05$<br>FC > 2 | Field | Martinelli et al., 2013 <sup>28</sup> |
| 17 | Valencia | 10 (5 healthy + 5 HLB), biologic replicates merged as one sample for each group | Mature fruit | – | SRP022979 | RNA-seq | DESeq | Adjusted $P < 0.05$<br>FC > 2 | Field | Martinelli et al., 2013 <sup>28</sup> |
| 18 | Hamlin | 4(2 healthy+2 HLB) | Dropped fruit | – | GSE101381 | RNA-seq | DESeq2 | Adjusted $P < 0.05$<br>FC > 2 | Field | Zhao et al., 2019 <sup>29</sup> |
| 19 | Hamlin | 4(2 healthy+2 HLB) | Retained fruit | – | GSE101381 | RNA-seq | DESeq2 | Adjusted $P < 0.05$<br>FC > 2 | Field | Zhao et al., 2019 <sup>29</sup> |
| 20 | Madam Vinous | 6 (3 healthy + 3 HLB) | Leaf | 7 weeks | Not available | RNA-seq | DESeq | Adjusted $P < 0.05$<br>FC > 2 | greenhouse | Yu et al., 2017 <sup>30</sup> |

**Table S3:** GO enrichment analysis of DEGs of *Citrus sinensis* in response to CLas infection based on nine different studies as specified in Table S2.

| GO_acc | Term type | Term | Query item | Bg item | p-value | FDR |
| --- | --- | --- | --- | --- | --- | --- |
| GO:0044464 | C | cell part | 880 | 1618 | 1.60E-49 | 3.20E-47 |
| GO:0005623 | C | cell | 880 | 1618 | 1.60E-49 | 3.20E-47 |
| GO:0044262 | P | cellular carbohydrate metabolic process | 92 | 111 | 2.70E-12 | 6.00E-09 |
| GO:0005506 | F | iron ion binding | 185 | 342 | 2.40E-10 | 3.70E-07 |
| GO:0003700 | F | transcription factor activity | 174 | 342 | 2.40E-08 | 1.80E-05 |
| GO:0003677 | F | DNA binding | 351 | 852 | 4.20E-07 | 0.00016 |
| GO:0003824 | F | catalytic activity | 2022 | 6096 | 4.20E-07 | 0.00016 |
| GO:0016491 | F | oxidoreductase activity | 492 | 1277 | 1.20E-06 | 0.00036 |
| GO:0043169 | F | cation binding | 529 | 1398 | 2.50E-06 | 0.00062 |
| GO:0046872 | F | metal ion binding | 524 | 1392 | 4.40E-06 | 0.00095 |
| GO:0008152 | P | metabolic process | 2064 | 6289 | 4.00E-06 | 0.0029 |
| GO:0009987 | P | cellular process | 1587 | 4729 | 3.80E-06 | 0.0029 |
| GO:0055114 | P | oxidation reduction | 443 | 1155 | 6.20E-06 | 0.0034 |
| GO:0016879 | F | ligase activity, forming carbon-nitrogen bonds | 37 | 47 | 1.90E-05 | 0.0036 |
| GO:0043565 | F | sequence-specific DNA binding | 91 | 174 | 2.30E-05 | 0.0039 |
| GO:0046914 | F | transition metal ion binding | 378 | 983 | 2.60E-05 | 0.004 |
| GO:0042221 | P | response to chemical stimulus | 84 | 159 | 3.50E-05 | 0.015 |
| GO:0020037 | F | heme binding | 165 | 396 | 0.00034 | 0.036 |
| GO:0016705 | F | oxidoreductase activity, acting on paired donors, with incorporation or reduction of molecular oxygen | 144 | 335 | 0.00028 | 0.036 |
| GO:0046906 | F | tetrapyrrole binding | 165 | 396 | 0.00034 | 0.036 |
| GO:0043167 | F | ion binding | 529 | 1481 | 0.00029 | 0.036 |
| GO:0005996 | P | monosaccharide metabolic process | 23 | 25 | 0.00015 | 0.045 |
| GO:0019318 | P | hexose metabolic process | 22 | 23 | 0.00015 | 0.045 |
| GO:0005976 | P | polysaccharide metabolic process | 47 | 77 | 0.00016 | 0.045 |

